# Prelimbic corticopontine neurons gate extinction learning

**DOI:** 10.1101/2024.08.14.607871

**Authors:** Hyoung-Ro Lee, Sung Hoon Choi, Yujin Kim, Seyeon Ko, Hyeji Kang, Inah Lee, Alan Jung Park, Suk-Ho Lee

## Abstract

The prelimbic cortex is essential for association of temporally separated events, and impedes extinction of learned association. The network mechanism underlying reversal learning, however, remains elusive. Here, we found that mitochondria-dependent post-tetanic potentiation at synapses onto prelimbic corticopontine neurons impedes extinction learning without affecting initial associative learning. Rats underwent trace fear conditioning followed by extinction sessions. Pharmacological inhibition of post-tetanic potentiation using tetaraphenylphosphnium (TPP), a mitochondrial Ca^2+^ release blocker, accelerated extinction of trace fear memory, leaving trace fear memory formation intact. Optogenetic inhibition of corticopontine, but not commissural, neurons phenocopied these effects. Electrophysiological recordings and Ca^2+^ imaging revealed that TPP treatment reduced the persistent activity of corticopontine neurons encoding tone presentation. Mechanistically, associative learning triggers the bursting activity of prelimbic pyramidal neurons that induces post-tetanic potentiation of corticopontine neurons, an effect blocked by TPP treatment. Thus, we identified a prelimbic cell type- and local circuit-specific mechanism that selectively gates extinction learning.

## Introduction

The prelimbic cortex is essential for learning associations between events that are temporally separated by a delay period, and impedes the extinction of previously learned associations^1,2^. A subpopulation of neurons in the prelimbic cortex persistently fire during the delay period^1,3^, which may convey information about the preceding event, thereby bridge events that are temporally separated. The information carried during a delay period is different, however, depending on the cell-types^3,4^. For example, intratelencephalic cells encode task-relevant information during a delay period of a working memory task, but pyramidal tract cells do not despite that they display robust persistent activity^4^. The role of persistent activity that is not directly relevant to a task performance is still elusive. One of possibilities is that persistent delivery of already learned information may hinder the modification of existing associations. Therefore, a circuit-specific approach to determine how persistent activity contributes to extinction learning will uncover the dynamic function of the prelimbic cortex.

Post-tetanic potentiation is a form of short-term plasticity where synaptic weight increases for tens of seconds after tetanic stimulation^5,6^. This phenomenon, by itself or by supporting persistent activity, can maintain task-relevant information during the delay period^7,8^. Prelimbic pyramidal neurons in layer 5 (L5) are categorized by their projection targets: commissural (COM) neurons that project to the contralateral prelimbic cortex, and corticopontine (CPn) neurons that project to the pontine nuclei^9^. We previously found that post-tetanic potentiation occurs selectively at two types of local synapses targeting prelimbic CPn cells: one is from L5 COM cells and the other from L2/3 pyramidal neurons^10^. Among these two types of synapses, only the latter (L2/3-CPn synapses) exhibited robust presynaptic mitochondrial Ca^2+^ release after tetanic stimulation, and thereby post-tetanic potentiation was induced. Notably, these forms of plasticity did not occur in other cortical areas including the infralimbic cortex^10^. The current understanding of prelimbic local circuitry, however, lacks a causal relationship between post-tetanic potentiation and persistent activity and their roles in associative learning and its extinction.

In this study, we tackled this issue using tetraphenylphophonium (TPP), which inhibits post-tetanic potentiation at prelimbic L2/3-CPn synapses by blocking presynaptic mitochondrial Ca^2+^ release with little adverse effects^6,10,11^. Convergent approaches combining in vivo electrophysiology, cell-type specific calcium imaging and optogenetic manipulations revealed that post-tetanic potentiation contributes to the persistent activity of prelimbic CPn neurons and interferes with extinction learning, but not associative learning.

## Results

### Inhibiting prelimbic post-tetanic potentiation accelerates trace fear memory extinction

Prelimbic persistent activity occurs during trace fear conditioning, and optogenetic silencing of the prelimbic cortex during the trace interval impairs trace fear memory acquisition^3,12^. To determine whether prelimbic post-tetanic potentiation contributes to persistent activity and associative learning, we examined the effects of TPP on prelimbic local network activity and trace fear conditioning.

We first conducted patch clamp experiments in rat brain slices to test whether TPP treatment affects the intrinsic electrical properties of prelimbic neurons. The bath application of 2 μM TPP did not exert significant impacts on input resistance, action potential (AP) threshold, and resting membrane potential of pyramidal neurons in layer 2/3, and 5, and of fast-spiking interneurons in layer 5 (**Fig. S1**). Next, we made slices in rats infused with 1 μl of 20 μM TPP into the prelimbic cortex. TPP treatment abolished post-tetanic potentiation at L2/3-CPn synapses within 300 μm, but not farther away, from the infusion site (**Fig. S2**).

We next examined the effect of TPP treatment on associative learning. Rats performed trace fear conditioning in which they learned to associate conditioned stimulus (CS, tone) and unconditioned stimulus (US, electric shock), traced by a 20 s interval. TPP (20 μM) or PBS (1 μl each) was injected into the prelimbic area 30 minutes before trace fear conditioning on day 1. Rats underwent five trials of CS-US pairing (**Fig. 1a**). Both groups displayed similar freezing levels during conditioning (**Fig. 1b**). On the following day, rats were presented ten times with the CS alone (tone test) to measure the formation and extinction of trace fear memory (**Fig. 1a**). The TPP group exhibited an accelerated extinction of conditioned fear. The freezing of the TPP group began to decrease after the 4th trial, whereas that of the PBS group kept high levels of freezing until the 6th trial (**Fig 1b**, **c**). It should be noted, however, that the freezing behavior of the TPP group was similar to that of the PBS group for the first three trials (**Fig 1b**, **c**), suggesting that the acquisition and retrieval of trace fear memory were not affected, but its extinction was accelerated by TPP infusion. Further, separate groups of rats were injected with either TPP or PBS 30 minutes before the tone test on day 2 (**Fig. S3a**). Similar to the results obtained when TPP was injected on day 1 (**Fig. 1b**), the freezing levels of the TPP group started to decrease earlier than those of the PBS group (**Fig. S3b**, **c**), whereas both groups showed similar freezing levels during conditioning on day 1 (**Fig. S3b**). Once TPP is absorbed into mitochondria by their negative potential, TPP is little washed out over time. Thus, TPP injected on day 1 may remain on day 2 and exert an inhibitory effect on mitochondria-dependent post-tetanic potentiation.

**Figure 1.**
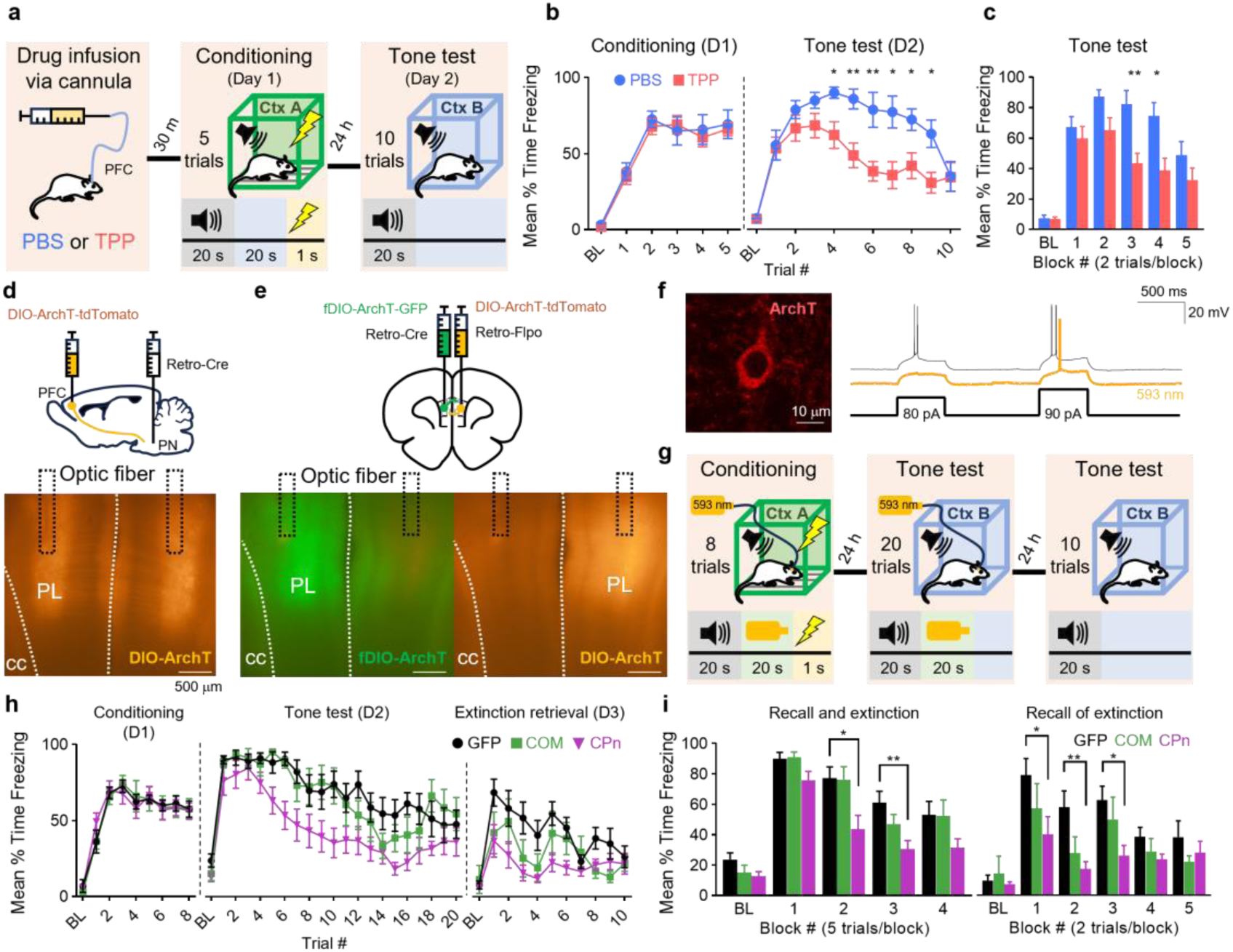
Both TPP infusion into the prelimbic (PL) area and opto-silencing corticopontine (CPn) neurons accelerated extinction of trace fear memory. **a-c:** Time course of freezing behavior after PBS or TPP injection into PFC 30 minutes before the trace fear conditioning. **a,** Schematic of experiment procedure. PBS or TPP was infused into the prelimbic cortex 30 minutes before the trace fear conditioning (TFC). **b,** Time course of freezing ratio of PBS or TPP-treated rats during trace fear conditioning (*left;* day 1) and tone test (*right*; day 2) (PBS, n = 6; TPP, n = 12; Conditioning on D1, F1,18 = 0.120, P = 0.733; Tone test on D2, F1,16 = 5.930, P = 0.027; Repeated measure two-way ANOVA (RM-ANOVA) for main effect of groups). **c,** Mean freezing ratios during tone test in PBS and TPP groups, presented as the average of 2 trials during tone test (F1,16 = 4.756, P = 0.044, RM-ANOVA for main effect of groups). **d-e:** Schematic of the CPn and COM neuron-specific expression of ArchT. **d,** *Upper*, to inactivate CPn activity, AAV2-retro-EF1a-Cre and AAV5-CAG-DIO-ArchT-tdTomato were injected into the pons and PFC, respectively. *Lower*, representative images of DIO-ArchT-tdTomato virus expression in CPn neurons and the optic fiber placement in the PL. Scale bar, 500 μm. **e,** *Upper*, to inactivate commissural (COM) neuron activity, the mixture of AAV2-retro-EF1a-Cre and AAV5-CAG-fDIO-ArchT-GFP and the mixture of AAV2-retro-EF1a-Flpo and AAV5-CAG-DIO-ArchT-tdTomato were injected into the contralateral and ipsilateral PFC, respectively. *Lower,* representative images of DIO-ArchT-tdTomato and fDIO-ArchT-GFP virus expression and optic fiber placement in the PL. Scale bar, 500 μm. **f,** *Left,* representative image of the ArchT-tdTomato-expressing PL neuron. Scale bar, 10 μm. *Right,* illumination of 593 nm light inhibited action potentials induced by step current injections (colored trace, at 80 and 90 pA) in the same cell. Scale bar, 500 ms, 20 mV. **g,** Schematic of neuron type-specific optical silencing experiment procedure. Illumination at 593 nm was applied during the trace interval of TFC and post-CS period of the tone test to inactivate CPn and COM neurons. **h,** Time courses of freezing ratio during TFC (*left*), tone test (*middle*), and extinction retrieval test (*right*) in control (GFP) and COM- or CPn-silenced animals (*black*, GFP; *green*, COM; *purple*, CPn) (For tone test, F2,27 = 9.960, P = 0.001, RM-ANOVA for main effect of group; for GFP vs CPn, P = 0.001; for GFP vs COM, P = 0.823, Tukey’s post hoc; For extinction retrieval test, F2,24 = 4.453, P = 0.023, RM-ANOVA for main effect of group; for GFP vs CPn, P = 0.018; for GFP vs COM, P = 0.283, Tukey’s post hoc). i, Mean freezing ratio presented as the average of 5 trials during the tone test (*left*, RM-ANOVA for main effect of group, F2,27 = 9.876, P = 0.001; Tukey’s post hoc for GFP vs CPn, P = 0.001; for GFP vs COM, P = 0.710), and as the average of 2 trials during the extinction retrieval tests (RM-ANOVA for main effect of group, F2,24 = 4.118, P = 0.029; Tukey’s post hoc for GFP vs CPn P = 0.023; for GFP vs COM, P = 0.333). Post hoc comparisons with Bonferroni correction were conducted to investigate differences between groups at each time point. All data are mean ± S.E.M., \**P* < 0.05, \*\**P* < 0.01, \*\*\**P* < 0.001.

To further test if TPP treatment has any effect on fear expression *per se*, we performed delayed fear conditioning, in which CS and US are temporally contiguous^13^ (**Fig. S3h**). The prelimbic cortex is required for fear expression in this task, while it is dispensable for conditioning^14^. The TPP group showed no significant difference from the PBS group in delayed fear acquisition and fear extinction (**Fig. S3i-j**). The prelimbic area is thought to hold task-relevant information during a trace period not only in trace fear conditioning but also in working memory tasks^1,4,15^. Given that TPP has little effect on CS-US association during trace fear conditioning, we performed an operant delayed-match-to-position working memory task to test whether TPP affects the ability to hold task-relevant information during a delay period (**Fig. S4a**). Rats treated with either TPP or PBS similarly performed in the working memory task (**Fig. S4b-d**), indicating that TPP has little effect on holding working memory. Overall, these findings suggest that blocking prelimbic post-tetanic potentiation at L2/3-CPn synapses accelerates extinction learning without affecting acquisition of trace fear memory.

### Silencing prelimbic corticopontine neurons accelerates fear memory extinction

Given that TPP treatment impairs post-tetanic potentiation at L2/3-CPn synapses (**Fig. S2**)^10^, we hypothesized that the TPP effects on extinction learning are mediated by preferential inhibition of CPn neurons. To test this, we performed cell type-specific optical silencing experiments. We injected AAV-DIO-ArchT-GFP or AAV-DIO-GFP into the prelimbic cortex, and retro-AAV-Cre into the pons or the contralateral prelimbic cortex to express the inhibitory opsin ArchT specifically in CPn and COM neurons, respectively (**Fig. 1d**, **e**). To express ArchT in COM neurons on the contralateral prelimbic cortex, we used a pair of fDIO-ArchT-GFP and retro-AAV-flipase (**Fig. 1e**). For the control group, AAV-DIO-GFP and retroAAV-Cre were injected into the prelimbic cortex and pons, respectively. As expected, CPn neurons were labeled only in L5, and COM neurons were observed both in layers 3 and 5 of the prelimbic cortex (**Fig. 1d**, **e**). We confirmed that ArchT-mediated hyperpolarizing photocurrents successfully inhibited the firing of the labeled neurons (**Fig. 1f**).

Rats were habituated to the conditioning chamber and presented with eight CS-US pairing trials for trace fear conditioning. We optogenetically silenced ArchT-expressing CPn or COM neurons during the trace interval of trace fear conditioning (day 1) and the post-CS period of the tone test. For the tone test, tone alone was presented 20 times in a distinct context (day 2) (**Fig. 1g**). Optogenetic silencing of either cell type did not affect freezing behavior during trace fear conditioning and the first five trial block of the tone test compared with the control group (**Fig. 1h-i**). On the other hand, silencing CPn, but not COM, neurons accelerated the extinction of conditioned fear, an effect similar to the TPP treatment (**Fig. 1h-i**). The day after extinction (day 3), the CPn-silenced group showed significantly lower freezing during the initial two trials compared with the other groups, further confirming that silencing CPn neurons facilitates extinction learning (**Fig. 1h-i**). Overall, these findings imply that TPP treatment accelerates extinction learning by inhibiting CPn neurons as silencing CPn neurons phenocopied TPP treatment.

### TPP inhibits persistent activity and accelerates extinction of prelimbic CS responses during tone tests

To directly measure the effect of TPP on prelimbic neuronal activity during extinction learning, we performed *in vivo* electrophysiological recordings (**Fig. 2a**). Behavioral procedures were the same as those for the optogenetic experiments (**Fig. 1g**). TPP or PBS was injected into the prelimbic cortex 30 min before trace fear conditioning on day 1. On the next day (day 2), we examined the activity of prelimbic neurons during the tone test (**Fig. 2a**). The CS responses of prelimbic neurons dissipated over trials, and the trial-dependent changes were accelerated by TPP (**Fig. 2b**), reflecting earlier decline of freezing behavior during the tone test in the TPP-treated and CPn-silenced groups (**Fig. 1b and 1h**). Based on z scores for firing frequencies and interspike intervals relative to their baseline values, neurons displaying distinctly high activity during 20 s tone presentation (CS period) of the first block (five trials/block) of the tone test were categorized into CS cells (see Methods). The time course of mean CS cell activity over 20 trials of the tone test is shown in **Fig. 2c**. When CS cell activity during the first half of CS period (CS-early phase) was analyzed, the activity of the TPP group was similar for the first five trials but decayed earlier than that of the PBS group (**Fig. 2d-e**). To examine the effects of TPP on the time course of CS cell activity before their CS responses decline, we compared their activity time courses averaged over the first block (trials 1-5) between two groups (**Fig. 2f**). Whereas two groups exhibited similar initial response to the tone presentation, CS cell activity in the TPP group decayed faster resulting in significantly lower activity during the second half of CS period (CS-late phase) and post-CS period (**Fig. 2g**). Accordingly, when we selected neurons exhibiting distinctly high activity during the post-CS period, the proportion of these persistently active (PA) cells among CS cells was lower in the TPP group (PBS, 26/60; TPP, 4/32; p = 0.0056), whereas the proportion of CS cells in the total number of cells was not different between two groups (PBS, 60/186; TPP, 31/124; p = 0.2122, **Fig. 2h**). Notably, similar effects were also observed when TPP was delivered 30 minutes before the extinction learning task (**Fig. S3d-g**). Collectively, these results indicate that TPP treatment selectively reduces CS-triggered persistent activity of prelimbic neurons without affecting early CS response. Together with the finding that TPP treatment accelerates extinction learning on day 2 (**Fig. 1**), these results suggest that prelimbic persistent activity during the tone test may play a role in maintaining CS-US associative memory, impeding extinction learning.

**Figure 2.**
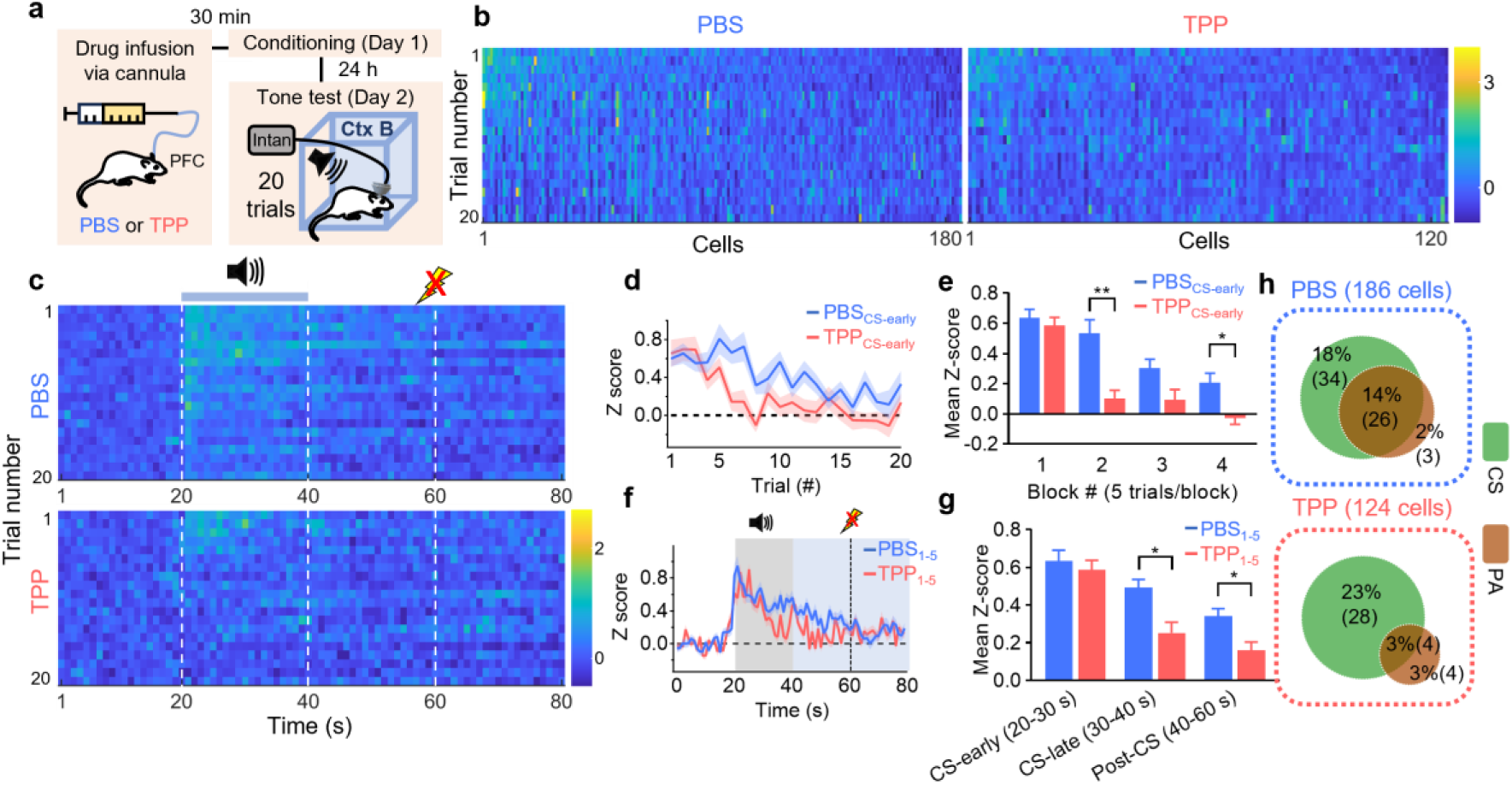
TPP infusion into PL reduces the persistent activity and accelerates extinction of prelimbic CS responses during tone test. **a,** Schematic of single unit recording during the tone test (Day 2) 30 min after PBS and TPP infusion into the PL area. **b,** Heatmap of z-scores of individual cells averaged over CS period of each trial (PBS, n= 186 cells from 10 rats; TPP, n=124 cells from 7 rats). Color bar, z-score. **c,** Heatmap of mean z-scores averaged over all CS-responsive cells of PBS (*upper*, n=60) and TPP groups (*lower*, n = 32) during the tone test. Vertical white lines indicate the CS period (20-40 s) and expected US timing (60 s). **d,** Activity changes of CS-responsive cells over trials in PBS (blue) and TPP (red) animals during the early 10 s of CS presentation (CS-early period). **e,** Mean z-scores of CS-responsive cells during CS-early period (20-30 s) across five trial blocks (5 trials/block). Linear mixed-effects (LME) models (TPP *vs*. PBS): F1,13 = 10.587, P = 0.006, for main effect of group; P = 0.0011 for trial block 2; P = 0.062 for trial block 3; P = 0.0436 for trial block 4; Tukey’s post hoc. **f,** Time-dependent changes of mean z-score of CS responsive cells averaged over first five trials in PBS (blue) and TPP (red) groups. Gray box, CS period (20-40 s). Vertical black dash line, expected US timing. **g,** Mean z-scores of CS cells averaged over three epochs shown in *f* (CS-early, CS-late, and post-CS periods). LME models for 1 s-binned activities between TPP vs. PBS groups: for CS-late period (30-40s), F1, 13 = 4.678, P = 0.0497; for post-CS period (40-60s), F1, 13 = 6.601, P = 0.0233. **h,** Proportions of CS and PA cells observed in the tone test. The numbers in the parentheses on the pie chart indicate cell counts. Chi-squared test: for CSPBS (60/186) vs. CSTPP (32/124), P = 0.275; for PAPBS (29/186) vs. PATPP (8/124), P = 0.0243. For LME models, time and drug treatment were set as fixed effects, while individual animals and neurons were set as random effects to account for repeated measures and clustering by animals. All data are mean ± S.E.M., Shaded areas represent S.E.M., *P < 0.05, ***P < 0.001.

### Early CS-responsive cells showing persistent activity belong to CPn cells

To specify the cell type whose activity is affected by TPP, it needs to classify tone-responsive cells into non-overlapping groups. To this end, we ordered tone-responsive cells (CS and PA cells) according to the timing of peak activity, and classified them into CS-early, CS-late and delay cells, which have a peak activity timing during early and late 10 s of CS presentation and post-CS period of the tone test, respectively (**Fig. 3a-c**). The proportions of these three cell types were not altered by TPP treatment (**Fig. 3c**). The time course of mean neuronal activity in each cell type, however, revealed that TPP selectively reduced the post-CS activity of CS-early cells (p = 0.0171, **Fig. 3d**), while it did not affect the activity of other cell types (**Fig. 3e-f**).

**Figure 3.**
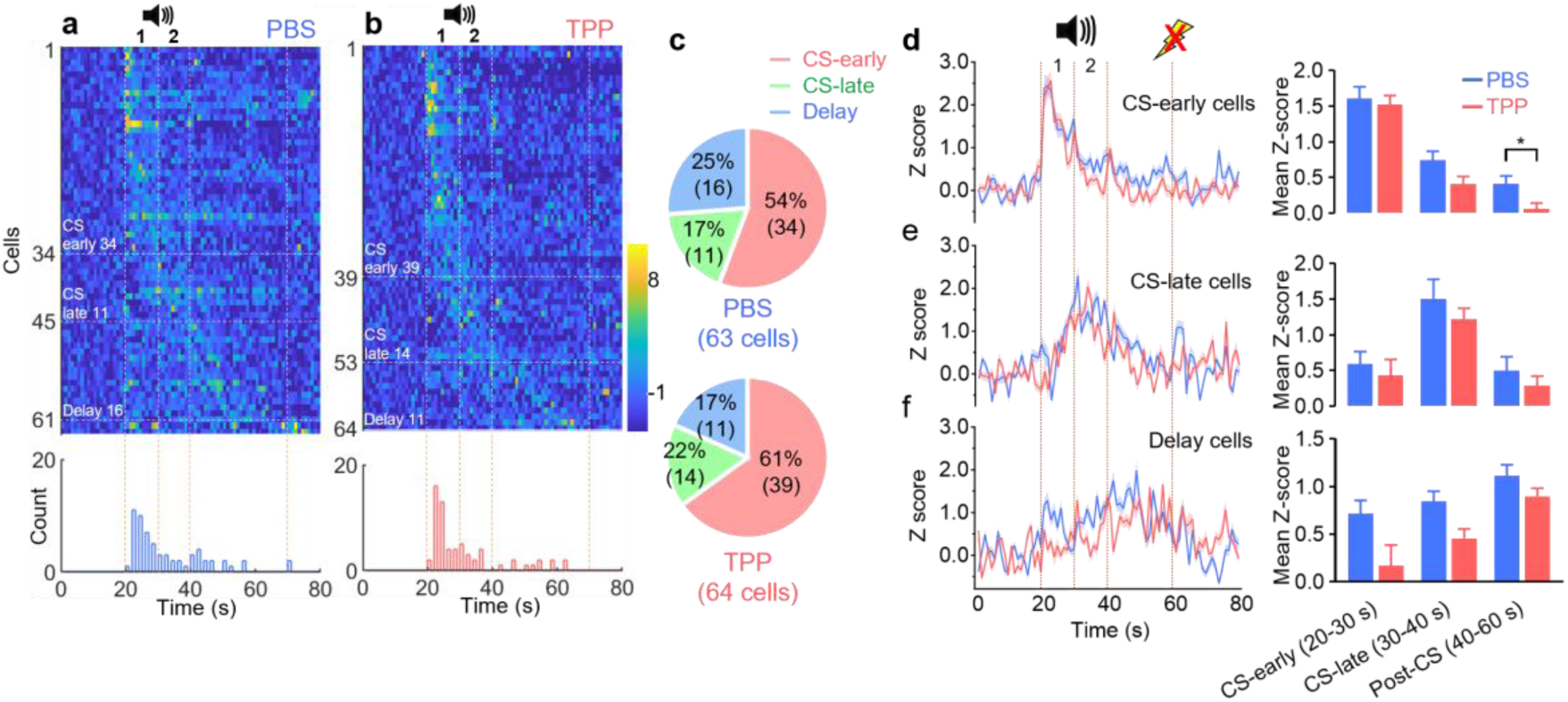
Effects of TPP on the activities of cell types classified by peak activity timing. **a-b,** Sequential activity of CS and PA cells in PBS and TPP injected groups. Data from experiments of day 1 and day 2 TPP injections (shown in Fig. 2 and Fig. S3, respectively) were merged for these plots to collect sufficient number of cells responsive to tone tests. *Upper,* Heat maps of activity time courses of individual cells (z-scores) averaged over first five trials of the tone test for PBS-(*a*) and TPP-(*b*) treated group. For each group, CS and PA cells were selected from electrophysiological recordings as shown in Fig. 2 and Fig. S3 from animals injected with PBS (*a*, n = 63 out of 186 total cells from 10 rats) and TPP (*b*, n = 64 out of 209 total cells from 12 rats), and sorted according to their timing of the peak z score. The peak timing in each cell was determined from a gaussian-filtered time sequence of z scores (σ = 4 s). *Lower,* Histograms for counts of cells displaying peak activity in each of 1 s bin. Vertical dashed lines demarcate the beginning of tone, first 10 s of CS period (CS-early), second 10 s of CS period (CS-late), and 10 s after expected US timing. **c,** Pie charts for the proportions of CS-early, CS-late and delay cells from PBS (upper, n = 63) and TPP (lower, n = 64) groups. CS and PA cells were categorized into the three types based on their peak activity timing. The numbers in the parentheses on the pie chart indicate cell counts. Chi-square test: for CS-early cell, P = 0.5387; for CS-late cell, P = 0.6874; for delay cell, P = 0.3609. **d-f,** Mean time-dependent activities of CS-early cells (*d*), CS-late cells (*e*), and delay cells (*f*) as shown in *a*-*b*. The cell numbers for the three cell types are shown in *c*. *Right,* Mean z-scores averaged over each of three epochs (CS-early, CS-late and post-CS period). LME models of 1 s-binned CS-early cell activities (PBS vs TPP): for CS-late period (30-40 s), P = 0.0709; for post-CS period (40-60 s), F1,16 = 7.076, P = 0.0171. LME models of delay cell activities (PBS vs TPP): for CS-early period, P = 0.698; for CS-late period, P = 0.241; for post-CS period, P = 0.6901. Time and drug treatment were set as fixed effects of LME models, while individual animals and neurons as random effects. All data are mean ± S.E.M., Shaded areas represent S.E.M., *P < 0.05.

Next, we examined cell type-specific Ca^2+^ spike activity during the tone test. The Ca^2+^ sensor GCaMP6f was specifically expressed in CPn and COM neurons using the same viral approaches described in **Fig. 1d-e** (**Fig. 4a-b**). The Ca^2+^ spike activity was recorded during the tone test on day 2 following trace fear conditioning, and analyzed using CNMF-E (**Fig. 4c-d**)^16^. The time course of CS cell activity averaged over first five trials was not different between CPn and COM cells (**Fig. 4e-f**). We selected tone-responsive cells (CS and PA cells), and further classified them according to their peak activity timing into CS-early, CS-late and delay cells from the heatmap of their sequential activity (**Fig. 4g-h**). COM cells had a significantly larger number of CS-early cells compared to CPn cells (p=0.001) (**Fig. 4g-i**). Time courses of activity averaged over each cell type are compared between COM and CPn cells in **Fig. 4j-l**. CS-early cells exhibited strong activity immediately following the CS onset both in COM and CPn cells, but the tone response of COM cells was rather transient compared to CPn cells (**Fig. 4j**). Notably, post-CS persistent activity of CS-early cells was observed specifically in CPn cells but not in COM cells (P = 0.0175, **Fig. 4j**), whereas post-CS activity in other cell types was not significantly different between COM and CPn cells (**Fig. 4k-l**). Because TPP specifically reduced persistent activity of CS-early cells (**Fig. 3a-b**), these findings suggest that the TPP effects on the CS-early cells are specific to CPn cells.

**Figure 4.**
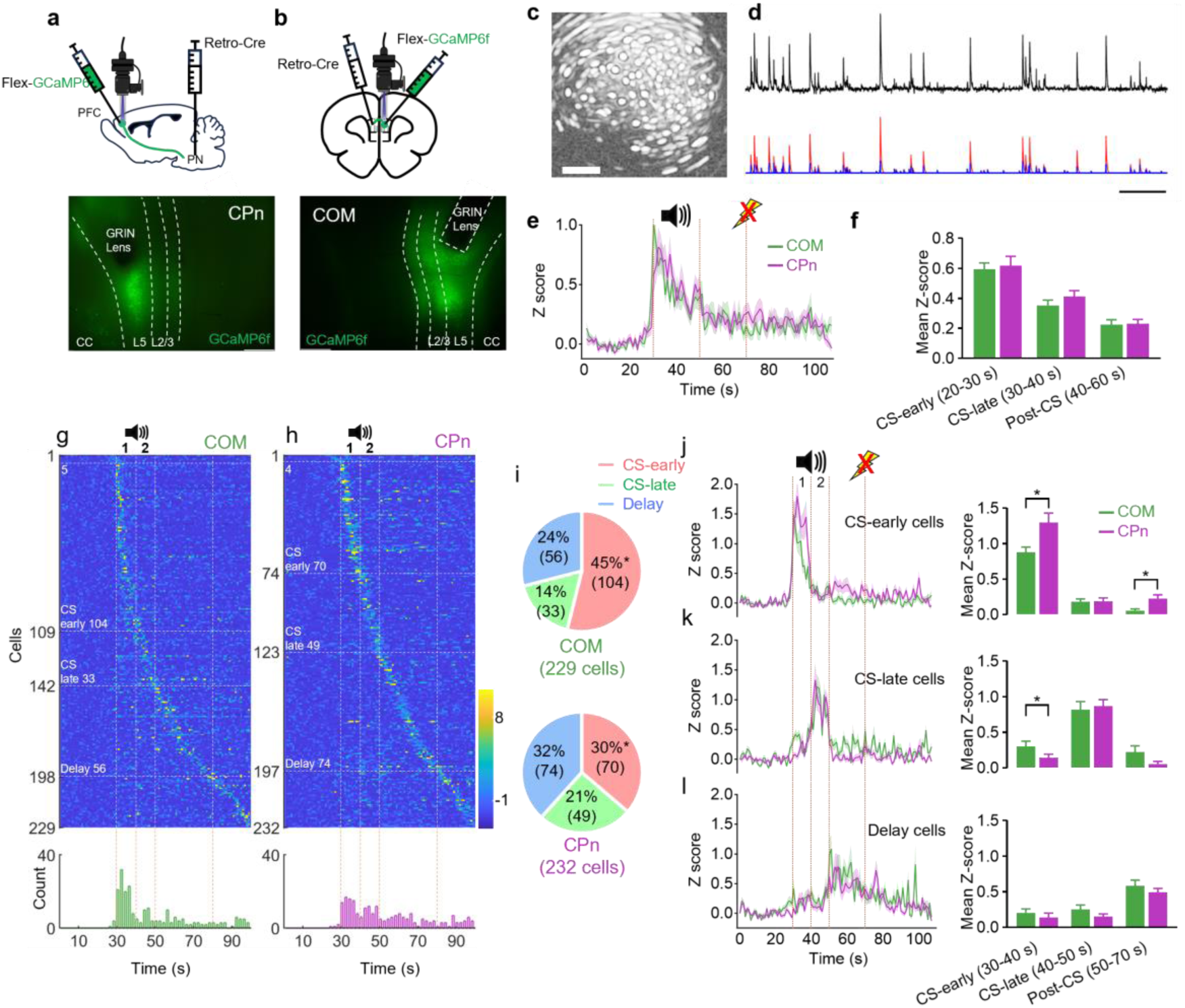
Cell-type specific activity during the tone test. **a-b,** Schematic of the methods for CPn (*a*) and COM (*b*) cell-specific virus expression. **a,** *Upper*, To record CPn activity, AAV2-retro-EF1a-Cre and AAV1-hsyn-flex-GCaMP6f were injected into the pons and the PL area, respectively. A GRIN lens was implanted into the PL. *Lower*, Representative image of GCaMP6f expression in the PL. **b,** *Upper*, To record COM activity, AAV2-retro-EF1a-Cre and AAV1-hsyn-flex-GCaMP6f were injected into the contralateral and ipsilateral PL, respectively. A GRIN lens was implanted into the ipsilateral PL. *Lower*, Representative image of GCaMP6f expression in the PL. Scale bar, 500 μm. **c,** Representative Miniscope image of COM cells obtained using a 1 mm GRIN lens. **d,** Representative raw and intermediate data used for CNMF-E analyses of Miniscope images. *Upper*, Raw calcium trace from an exemplar neuron. *Lower*, Simulated calcium trace (red) calculated from convolution of inferred spike (blue) and a unitary calcium transient, which were estimated from CNMF-E. Scale bar, 1 min. **e,** Mean z-score of all CS responsive cells for COM (green; n = 192 out of 618 total cells from 13 rats) and CPn (purple; n = 186 out of 682 total cells from 13 rats) neurons. For each cell, z scores were averaged over first five trials of tone test. **f,** Time-averaged z-scores of CS cells for three epochs (CS-early, CS-late, and post-CS periods). **g-h,** *Upper*, Heat map was plotted similar to Fig. 3a-b for COM (*g*) and CPn (*h*) cells (COM cells: n = 229 out of 618 total cells. CPn cells: n = 232 out of 618 total cells). For each cell, time series of 1 s binned z scores were averaged over first five trials of the tone test. *Lower,* Histogram for counts of cells displaying peak activity in each 1 s bin. Vertical dashed lines demarcate the beginning of tone, first 10 s of CS period (CS-early), second 10 s of CS period (CS-late), and 10 s after expected US timing. **i,** Pie charts for proportions CS-early, CS-late and delay cells, into which CS and PA cells of COM (upper) and CPn (lower) cells were categorized according to their peak timing. The numbers in the parentheses on the pie chart indicate cell counts. Chi-square test (COM vs CPn): for CS-early cell, P = 0.001; for CS-late cell, P = 0.0781; for delay cell, P = 0.0945. **j-l,** *Left*, Mean time-dependent activities (z scores) of CS-early cells (*j*), CS-late cells (*k*) and delay cells (*l*) of COM (green) and CPn (purple) cells, shown in *g* and *h*. The numbers of CS-early, CS-late and delay cells are shown in *j*. *Right*, Mean z-scores averaged over each of three epochs (CS-early, CS-late and post-CS period) for CS-early (j), CS-late (k) and delay (l) cells. Green, COM cells. Purple, CPn cells. LME models for 1 s-binned CS-early cell activities between COM vs. CPn: CS-early period (30-40 s), F1,24 = 5.5889, P = 0.0265; post-CS period (50-70 s), F1, 24 = 6.5147, P = 0.0175. LME model of CS-late cell activities during CS-early period (COM vs CPn), F1,22 = 4.4251, P = 0.0471; post-CS period (50-70 s), F1, 22 = 4.101, P = 0.0551. Time and cell types (COM vs CPn) were set as fixed effects of LME model, while individual animals and cells were set as random effects. All data are mean ± S.E.M., Shaded areas represent S.E.M., *P < 0.05.

### The firing pattern of a L2/3 pyramidal neuron induces post-tetanic potentiation in L5 CPn neurons

We previously showed that TPP inhibits post-tetanic potentiation evoked by 25 Hz stimulation at prelimbic L2/3-CPn synapses *in vitro*^10^, but it remains to be tested whether CS-evoked L2/3 pyramidal neuronal activity *in vivo* can induce post-tetanic potentiation in CPn neurons. We recorded L2/3 neuronal activity patterns during first five trials of the tone test (**Fig. 5a**). The activity time course and proportions of CS and PA cells in L2/3 neurons were not different from those of layer-nonspecific recordings (**Fig. 5b-d**). The mean firing rate of CS cells in L2/3 increased from 1.5 Hz to 2.7 ± 0.3 Hz in response to CS (**Fig. 5e-f**). The distribution of inter-spike intervals (ISIs) averaged over CS cells revealed that ISIs shorter than 120 ms significantly increased in the CS period compared to the baseline (**Fig. 5g**), indicating that burst firings are responsible for the CS-evoked increased activity. We used the firing pattern of an exemplar CS cell during tone presentation as a stimulation template (**Fig. 5h-l**), and tested whether this patterned stimulation of L2/3 induces post-tetanic potentiation in CPn cells (**Fig. 5h-m**). To mimic in vivo conditions, recordings were done at 32℃ and 1.3 mM [Ca^2+^] in the bath. Under these conditions, post-tetanic potentiation was readily induced at L2/3-CPn synapses (**Fig. 5n-p**), and significantly reduced by bath-applying TPP (**Fig. 5n-p**), suggesting that CS-evoked L2/3 prelimbic activity can induce TPP-sensitive post-tetanic potentiation at L2/3-CPn synapses.

**Figure 5.**
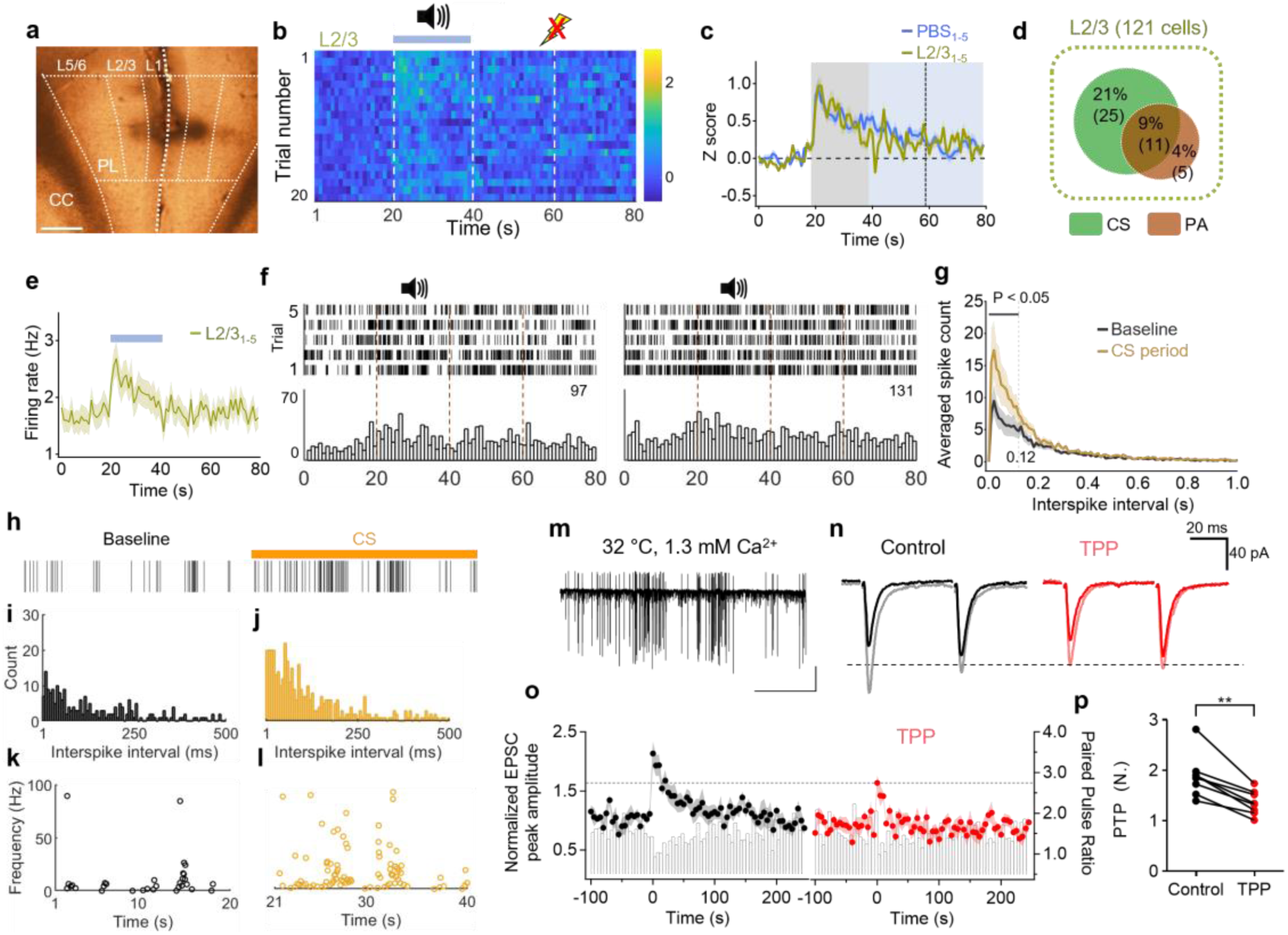
In vivo activities of prelimbic L2/3 pyramidal neurons can induce TPP-sensitive post-tetanic potentiation in L5 CPn neurons. **a,** Representative image for the location of streotrodes bundles to record prelimbic L2/3 neuron activity. Scale bar, 500 μm. **b,** Heatmap of z-scores averaged over all L2/3 CS responsive cells (n = 36/121 cells from 4 rats) during tone test. Vertical white lines, CS period and expected US timing. **c,** Time-dependent changes of mean z-score of L2/3 CS responsive cells averaged over first five trials of the tone test. For comparison, the z-score trace from layer-nonspecific recordings (denoted as PBS) is reproduced from Fig. 2f. Gray box, CS period. Vertical dashed line, expected US timing. **d,** Percentages of CS and PA cells in prelimbic L2/3 during tone test (CSL2/3, 36/121; PAL2/3, 16/121). The numbers on the pie chart represent percentages outside the parentheses and cell counts inside the parentheses. **e,** Time-dependent changes of mean spike frequency of all CS responsive cells in L2/3 (n = 36/121). Horizontal bar, CS period. **f,** Spike raster plots (*upper*) and corresponding post-event time histograms (PETHs) of spike counts (*lower*) of two representative L2/3 CS cells for 1-5 trials. Bin width of PETH, 1s. **g,** Mean spike counts of L2/3 cells as a function of inter-spike intervals (ISIs) during the baseline (black) and CS periods (orange). Bin width, 0.01 s. For each CS cell, spike counts were summed over 20 s of baseline and CS period over first five trials. For visibility, spike counts with ISI > 1 s were truncated. For ISIs shorter than 120 ms of baseline vs. CS period, F1, 6965 = 21.631, P < 0.05, LME model. Gray vertical dashed line indicates ISI at 120 ms. **h,** Spike activity of the cell #97 shown in *f* (*left*) during baseline and CS period in the 1st trial. **i-j,** Histogram of ISIs of cell #97 in the baseline and the CS period of 1-5 trials. Note that most ISIs were shorter than 200 ms for both periods. **k-l,** Plots of instantaneous frequencies (1/ISIs) of cell #97 during baseline and the CS period of the 1st trial. **m,** EPSCs trace at L2/3-CPn synapses evoked by patterned stimulation mimicking the firing pattern of *h* during CS period. **n,** Representative paired-pulse EPSCs before (*left*; black) and after TPP (*right*; red) applications. Dotted line indicates the peak EPSC amplitude after patterned stimulation under the TPP conditions. **o,** Time courses of mean baseline-normalized EPSC amplitudes at L2/3-CPn synapses before and after patterned stimulation applied at t = 0. Experiments were done in the conditions similar to in vivo (1.3 mM [Ca^2+^]o, 32 °C). Left, control (n = 8). Right, after TPP application (n = 8). **p,** Post-tetanic potentiation at L2/3-CPn synapses before and after applying TPP. For PTP of control vs TPP, P = 0.0078, Wilcoxon matched pairs signed rank test. All data are mean ± S.E.M., Shaded areas represent S.E.M., **P < 0.01.

### TPP also reduces persistent activity during trace fear conditioning

Because TPP inhibited persistent activity during the tone test, we further examined whether TPP has similar effects during trace fear conditioning on day 1 (**Fig. 6a**). Rats were habituated to the conditioning chamber and presented with three trials of CS alone (trial 1-3). Then, the rats underwent subsequent eight CS-US pairing trials (trial 4-11) for trace fear conditioning. The CS-responsive activity emerged in the trial 5, whereas CS presentation alone (trial 1-3) did not alter prelimbic activity relative to baseline (**Fig. 6b-d**). The effects of TPP on the network activity during trace fear conditioning were similar to those during the tone test. TPP treatment reduced the persistent activity of prelimbic neurons without affecting early CS responses (**Fig. 6e-f**). The proportion of PA cells among CS cells were lowered by TPP (**Fig. 6g**), whereas that of CS cells was not altered by TPP (59% vs. 55%), indicating that TPP specifically reduces persistent activity of prelimbic neurons not only in tone test but also in trace fear conditioning.

**Figure 6.**
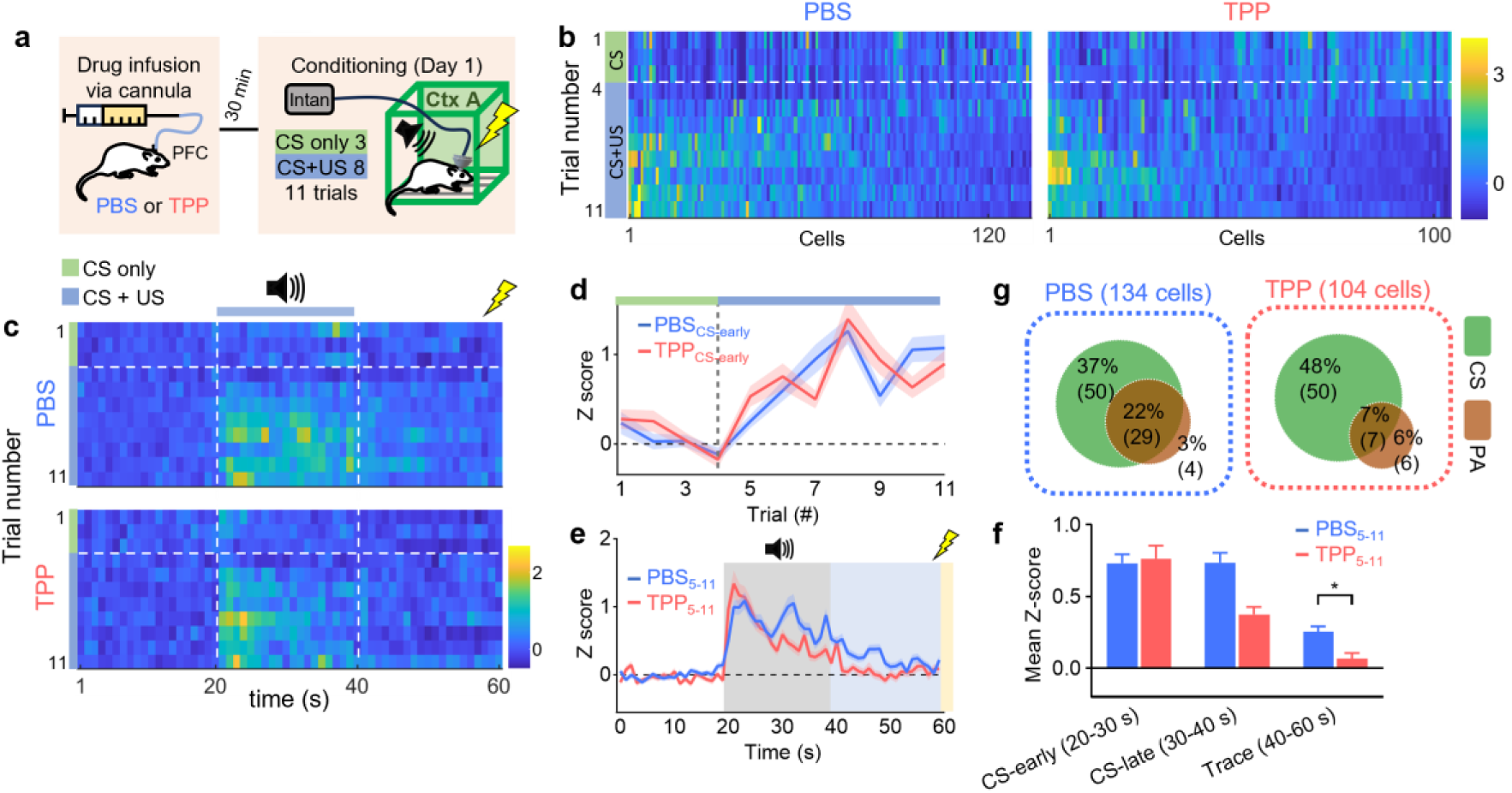
TPP infusion into the mPFC reduces the persistent activity of PL neurons during TFC. **a,** Schematic of single unit recording during TFC 30 min after PBS and TPP injection into the prelimbic cortex. Rats were habituated to the conditioning chamber and presented with three trials of CS alone (trial 1-3), and then underwent subsequent eight CS-US pairing trials (trial 4-11) for trace fear conditioning. **b,** Heatmap of z-scores of individual cells were averaged over the CS period of each trial (*left*, PBS, n= 134 cells from 5 rats; *right,* TPP, n=104 cells from 5 rats). Color bar, z-score. Cells were sorted based on their activities. **c,** Heatmap of z-scores averaged over all CS-responsive cells of PBS (*upper,* n = 79) and TPP groups (*lower,* n = 57) as a function of time. CS-only and CS+US trials are demarcated by the horizontal white line. The vertical white lines divide pre-CS, CS and trace periods. Color bar, z-score. **d,** Mean activity changes of CS-responsive cells over trials in PBS (blue) and TPP (red) animals during the CS-early periods. **e,** Mean z-scores of CS-responsive cells as a function of time. Z-scores shown in *c* were averaged over trials 5 to 11 for the PBS and TPP groups. The CS period is indicated by a gray box, the trace interval by a blue box, and the US presentation by a yellow box. **f,** Summary for the mean z-scores averaged over three epochs shown in *e* (CS-early, CS-late, and trace periods). For 1 s-binned activities during the trace period of TPP *vs* PBS group, F1,8 = 5.846, P = 0.041, LME model with time and drug treatment set as fixed effects, and individual animals and neurons as a random effect to account for repeated measures and clustering by animals. **g,** Percentages of CS and PA cells observed during conditioning. The proportion of PA cells among CS cells were significantly lower in the TPP group than the PBS group (29/79 vs. 7/57, p = 0.0028), whereas that of CS cells was not altered by TPP (59% vs. 55%). Chi-squared test: for CSPBS (79/134) vs. CSTPP (57/104), P = 0.191; for PAPBS (33/134) vs. PATPP (13/104), P = 0.0120. All data are mean ± S.E.M., Shaded areas represent S.E.M. *, P < 0.05.

## Discussion

Post-tetanic potentiation at the prelimbic L2/3-CPn synapse is induced by post-tetanic mitochondrial Ca2+ release via mitochondrial Na^+^/Ca^2+^ exchanger, which is blocked by TPP^10^. In the present study we examined the network and behavioral consequences of inhibition of post-tetanic potentiation at these synapses using TPP. TPP not only accelerated extinction learning but also selectively reduced persistent activity of CS-encoding CPn neurons during retrieval of trace fear memory. This notion is supported by *in vivo* electrophysiological recordings and Ca^2+^ imaging, which revealed that persistent activity of early CS-responsive cells (CS-early cells) was observed among CPn cells but not among COM cells, and it was selectively reduced by TPP. Moreover, optogenetic silencing of CPn cells during post-CS period phenocopied the behavioral effects of TPP. Finally, CS-evoked characteristic firing patterns of perlimbic L2/3 pyramidal neurons successfully induced TPP-sensitive post-tetanic potentiation at L2/3-corticopontine synapses. Together with our previous study^10^, these results suggest that post-tetanic potentiation at prelimbic L2/3-CPn synapses maintains persistent activity of prelimbic CPn neurons during retrieval of trace fear memory, which may impede extinction learning.

The present study relies on the pharmacological agent TPP, which blocks mitochondrial Na^+^/Ca^2+^ exchanger, the primary pathway of post-tetanic mitochondrial Ca^2+^ release^6,17,18^. Previous studies indicate that TPP specifically inhibits mitochondria-dependent post-tetanic potentiation with little adverse effects. Low concentration TPP did not affect presynaptic AP shapes, Ca^2+^ current, mitochondrial respiration and short-term plasticity at < 50 Hz *in vitro* ^11,17,19^. In line with these findings, the present study showed that TPP has little effect on the intrinsic properties including after-depolarization, firing frequency-current injection relationship, Input resistance, resting membrane potential and AP threshold in L2/3 and L5 pyramidal cells and fast-spiking interneurons (**Fig. S1**). The present study showed that TPP did not affect the early network response to CS (**Fig. 2f, 3d, 6e**), but it specifically reduced the delayed response, arguing against a possibility that TPP has non-specific adverse effects. Moreover, our behavioral tests showed that TPP effect was specific to extinction learning with little effects on trace fear conditioning and the working memory task, supporting little adverse effects of TPP. Nevertheless, we cannot rule out the adverse effects of TPP treatment, and further examinations should follow.

Inactivation of the prelimbic cortex during trace intervals impaired trace fear memory formation, suggesting that prelimbic activity during the trace interval contributes to temporal associative learning^3,20^. In our study, however, TPP did not affect fear memory formation even though it significantly reduced persistent activity during the trace interval. This finding was supported by little effect of TPP on the working memory task (**Fig. S4**), suggesting that TPP does not impair network activity that links temporally separated CS and US. It is possible that TPP-insensitive persistent activity may convey CS information, given that TPP has little effects on persistent activity of CS-late and delay cells (**Fig. 3**). Alternatively, a change in the population activity of the prelimic network rather than persistent activity itself may carry the CS information during trace intervals.

Activity and plastic changes in the prelimbic area are required for new memory formation of CS-US association during TFC^3,13,21^, but not for retrieval of trace fear memory in already conditioned animals^2^. Nevertheless, extinction of trace fear memory was accelerated by inhibition of the prelimbic cortex^2^. More generally, silencing of the prelimbic cortex promoted reversal learning in an operant conditioning task of cue-reward pairing while it did not affect recall of already learned rule^2^. Our results elaborated these previous findings by elucidating prelimbic cell type-specific activity that impedes extinction learning. We showed that acceleration of extinction learning closely correlates with TPP-induced inhibition of prelimbic persistent activity during the post-CS period of the tone test. As aforementioned, we presented evidence supporting that TPP-sensitive persistent activity occurs in CPn cells. Therefore, we conclude that post-CS persistent activity of prelimbic CPn cells interferes with extinction learning. It remains to be elucidated whether prelimbic persistent activity also contributes for an animal to adhere to the previously learned rule even if the animal is repeatedly exposed to an alternative rule.

Our data indicate that CS presentation induces burst firing of prelimbic L2/3 pyramidal neurons, which can trigger post-tetanic potentiation in CPn neurons thereby interfering with rapid extinction learning. The mediodorsal thalamus may drive this bursting activity because it comprises the major long-range inputs and burst firings of thalamic afferents provide net excitatory inputs to prelimbic L2/3 pyramidal neurons^22,23^. Indeed, enhanced burst firing in the mediodorsal thalamus impairs extinction learning^24^. Investigating direct interactions between the prelimbic cortex and mediodorsal thalamus during extinction learning will confirm this notion.

Post-tetanic mitochondrial Ca^2+^ release is inhibited by mild mitochondrial depolarization, which observed in non-symptomatic stage of Alzheimer disease^25^. Consistently we have showed that young adult Alzheimer disease model mice phenocopied the effects of TPP: mild mitochondrial depolarization in L2/3 pyramidal neurons and the lack of post-tetanic potentiation at L2/3-CPn synapses, phenocopying the effect of TPP treatment^10^. Therefore, TPP-induced network and behavioral changes would have great implications in preclinical signs of brain disorders involving mitochondrial dysfunction.

## Materials and Methods

### Animals

Sprague Dawley rats (21-42 days) were used for the main study. C57BL/6J mice (16-18 weeks) were used in working memory experiments. Animals were housed under a 12-h light/12-h dark cycle (light from 8 a.m. to 8 p.m.) with constant room temperature of 21 °C and 65% humidity. For trace fear conditioning using rats, food and water were available ad libitum before the start of experiments. For working memory tasks using mice, food was restricted such that the body weight is reduced to 85-90% of standard body weight of the same aged mice (typically, 23 - 24 g). All experiments were performed during the light cycle. All experimental procedures were approved by Seoul National University Institutional Animal Care and Use Committee (SNU-200831-3-3, SNU-090115-7-9).

### Stereotaxic surgery

#### Cannula implantation surgery

For infusion of TPP or PBS into the PL, a set of bilateral guide cannulae, dummy, and caps were implanted into the PL area (guide cannula: 26 G, center-to-center distance: 1.2 mm for rat and 1 mm for mouse; dummy needle, 33G; PlasticsOne, USA). The stereotaxic coordinates were as follows: For rat PL: AP 2.3, ML ±0.6, DV -3.4; for mouse PL: AP 1.8, ML ±0.5, DV -1.6. Representative images and locations of the bilateral infusion cannulae used in present study are illustrated in **Fig. S5**. After implantation, the cannulae assemblies were fixed to the skull with dental cement (Poly-F plus, Dentsply Sirona, USA). After surgery, the animals recovered in the infrared lamp. Two weeks later, PBS or TPP were injected using a Hamilton syringe and infusion pump (volume, 1 μl/side; speed, 100 nl/min for 10 minutes). The injection cannula (33G, PlasticsOne, USA) was connected to the Hamilton syringe via polyethylene tubing (PE50, BD Intramedic, USA). The infusion cannula remained for 5 minutes to allow diffusion of the drug. Trace fear conditioning or tone test were performed 30 minutes after the injection was finished.

#### Cell-type specific optogenetic inactivation surgery

For specific inactivation of CPn neurons, AAV2-retro-EF1a-Cre (Addgene #55636, 1 × 10^13^ genomic copies (gc)/mL) and AAV5-CAG-DIO-ArchT-tdTomato (Addgene #28305, 1 × 10^12^ gc/mL) were bilaterally injected into the pons and PL, respectively. To inactivate COM cells, the mixture of AAV2-retro-EF1a-Cre (2 × 10^13^ gc/mL) and AAV5-CAG-fDIO-ArchT-GFP (plasmid: Addgene #124640; virus: made in KIST, 2 × 10^13^ vg/mL) and the mixture of AAV2-retro-EF1a-Flpo (Addgene #55637, 2 × 10^13^ vg/mL) and AAV5-CAG-DIO-ArchT-tdTomato (2 × 10^12^ gc/mL) were injected into the contralateral and ipsilateral PFC, respectively. For GFP expression in CPn neurons as a control group, AAV2-retro-EF1a-Cre (1 × 10^13^ gc/mL) and AAV5-CAG-DIO-EGFP-WPRE (Addgene #51502 1 × 10^13^ gc/mL) were bilaterally injected into the pons and the PL, respectively. Animals were anesthetized with 5% (v/v) isoflurane during surgery. The scalp was shaved, and the animal was placed into the stereotaxic apparatus (Stoelting, USA). The scalp was disinfected with 70% (v/v) alcohol and Povidone-Iodine, and craniotomy was performed. The stereotaxic coordinates were as follows: For pons, AP -7, ML ±1, DV -10; For PL, AP 2.3, ML ± 0.6, DV -3.4. Virus was injected using an Ultra Micro Pump (UMP3, WPI, USA) connected to a microcontroller (MICRO2T, WPI, USA). The virus was loaded into a Hamilton syringe (7653-01, Hamilton, USA) and a needle (7762-08, 28 G, Hamilton, USA). The virus solution of 1 μl was injected at a rate of 100 nl/min for 10 minutes. The injection syringe remained for 5 minutes to allow diffusion of the virus. Immediately after surgery, the animals were recovered under the infrared lamp. Two weeks after virus injection, optic fibers (R-FOC-L200C-39NA, 1.25 mm ceramic ferrule, 200 um Core, 3.5 mm length, 0.39 NA, RWD, China) were bilaterally implanted into the PL for 593 nm laser illumination. Behavioral experiments began two weeks after optic fiber implantation.

#### Shuttle drive surgery

An in vivo electrode drive (called shuttle drive) was built according to Voigts et al (2020) with modifications^26^. The drive was constructed by combining a bundle of 16 stereotrodes, a 3D-printed body, and an electrode interface board (EIB). Stereotrodes were made by folding, twisting, and heating the Formvar-coated nichrome wire (761000, A-M systems, Netherlands). An electrode interface board (EIB) consisted of a connector (A79026-001, Omnetics, USA) and an open-source printed circuit board (PCB, SCITECH, Korea). Individual stereotrode wires were connected to the EIB with gold pins (OPES-7013, Open Ephys, Portugal). For the TPP infusion experiment, a 26-gauge stainless tube was passed through the hole in the middle of the EIB and the center of the 3D-printed body (**Fig. S6a**). The EIB was attached to the 3D-printed body and then glued with epoxy. A 28-gauge silver coated copper wires (TAWG28, SME, Korea) soldered with skull screws were connected to the ground and reference holes of the EIB using gold pins. Before the surgery, a bundle of stereotrodes was cut with sharp scissors, and was gold-plated to lower the impedance to 200–300 kΩ.

Rats were anesthetized with 5% (v/v) isoflurane. After the hair was removed, rats were moved into a stereotaxic apparatus (Stoelting, USA). the scalp was disinfected with 70% (v/v) alcohol and Povidone-Iodine. After the scalp was incised, the skull surface was cleaned using H2O2 and the craniotomy site was marked (PFC AP 2.3, ML 1.6, DV -3.5, Angle 16°). Then, a circular 1-mm-diameter craniotomy was made. A stereotrodes bundle installed in a hand-made shuttle drive was slowly lowered to a depth of 3.5 mm. A ground screw was placed on the skull of cerebellar area and a reference screw was placed in the skull of the olfactory bulb area. The surface of exposed skull was covered with a dental cement (Super-Bond C&B, Sun Medical, Japan). On the top of it, the shuttle drive was fixed using another dental cement (Poly-F plus, Dentsply Sirona, USA). Immediately after surgery, the animals recovered under the infrared lamp.

#### Cell type specific in vivo calcium imaging surgery

GRIN lens implantation was performed in two alternative ways depending on the use of silk fibroin. 1) AAV2-Cre (1 × 10^13^ gc/mL) was injected into the pons or contralateral PFC, and AAV-flex-GCaMP6f (Addgene #100833, 2 × 10^13^ gc/mL) was injected into the PL; One week later, a 1 mm or 0.6 mm GRIN lens was implanted in the PL. 2) AAV2-Cre (1 × 10^13^ gc/mL) was injected into the pons or contralateral PFC; One week later, after applying AAV-flex-GCaMP6f (2 × 10^13^ gc/mL) virus to the surface of a GRIN lens using silk fibroin, the lens was implanted in the PFC. The virus was injected for 10 minutes using a Hamilton syringe and an Ultra Micro Pump with a Micro Controller (volume 1ul, speed 100 nl/min). A week after the virus injection, a GRIN lens was implanted. For the first method, the lens was sterilized with 70% ethanol and implanted. For the second method, the lens was sterilized with UV light. Then, the virus was applied to the surface of the lens distal tip using silk fibroin (Merck, USA). After mixing AAV-flex-GCaMP6f (2 × 10^13^ gc/ml) and silk fibroin in a 1:1 ratio, 1 μl of the mixture was applied to the surface of the lens and dried at room temperature for at least 1 hour. The animals were anesthetized and fixed in a stereotaxic apparatus again and a circular 1-mm-diameter craniotomy was made. Superficial cortical tissue was gently aspirated with the tip of a 22-gauge blunted needle connected to a vacuum source. After this, a 1.0 mm or 0.6 mm-diameter GRIN lens (Inscopix, USA) was slowly lowered to the depth of 3.5 mm while constantly washing the craniotomy site with sterile PBS. The surface of the skull was covered with dental cement (Super-Bond C&B, Sun Medical, Japan). Then, the GRIN lens was fixed on the head using dental cement (Poly-F plus, Dentsply Sirona, USA). Immediately after surgery, the animals recovered under the infrared lamp. To avoid surgery-related tissue swelling, rats were injected with dexamethasone (1 mg/kg) for one week after surgery.

### Behavior

#### Trace Fear conditioning

The fear conditioning was performed in a context A, which is a rat conditioning cage (H10-11R-TC, Coulbourn Instruments, USA) located inside a sound-attenuating box (SCITECH, Korea) with an LED lamp. The auditory cue was presented via a speaker module (H12-01R, Coulbourn Instruments, USA) located on one side of the conditioning chamber. The floor of the conditioning cage consisted of stainless-steel bars to deliver a footshock (H10-11R-TC-SF; Coulbourn Instruments, USA). Two types of context B were used for the tone test depending on the type of experiments. One is for behavior tests after TPP or PBS infusion, in which the freezing behavior was monitored (Context B). The other is for in vivo recordings and optogenetic inactivation experiments, in which in vivo recording of neuronal activities or laser illumination was done in parallel with observing freezing behaviors (Context B’). Context B used for the behavior test was hexagonal (32 × 34 × 50 cm) and consisted of a white acrylic floor and wall located in the floor of the isolated room. The auditory cue was delivered through speakers located on either side of Context B. Context B’ used for in vivo recording and cell type specific inactivation experiments was rectangular (36 × 21x 25 cm), consisted of a black plastic floor and glass wall with blue paper on the outside, and was located inside a sound-attenuating box. To quantify freezing behaviors, a python-based open source code ezTrack^27^ was used. The parameters we used for this analysis were a motion cutoff of 10 - 25, a freezing threshold of 30, and a minimum freeze duration of 30 samples. The behavior was recorded with a CCD camera.

For TPP-infusion experiments, rats were subject to fear conditioning two weeks after cannula implantation. TPP or PBS was delivered to the PL area using bilateral infusion cannulae, each of which was connected to a Hamilton syringe. TPP or PBS was infused 30 minutes before the conditioning (**Fig. 1a**) or tone test (**Fig. S3a**). In trace fear conditioning (day 1), the rats received five CS-US pairings with pseudo-random intertrial intervals between 160 and 200 s. The CS-US pairing consisted of tones (2.5kHz, 20 s, 75 dB) and a footshock (0.5 mW, 1 s). CS and US are separated by a 20 s trace interval. In DFC, rats received five CS-US pairings (**Fig. S3h**). CS-US pairing of DFC consisted of tone presentations (2.5kHz, 20 s, 75 dB) that coterminated with footshocks (0.5 mA, 1 s). The intertrial intervals were 180 ± 20 s. The next day, in the tone test (day 2), the rats received ten presentations of tones (20 s, 2.5 kHz, 75 dB) in context B. The intertrial intervals of tone presentations were 90 ± 20 s.

For cell type specific optogenetic inactivation experiments (**Fig. 1g**), the rats were subject to trace fear conditioning two weeks after optic fiber implantation. To illuminate the PL area with 593 nm laser during behavioral experiments, the optic fibers implanted in the rat’s head were connected to the 593.5 nm laser (YL593T6-050FC, Lasercentury, China) before behavior tests via a fiber optic patch cable (R-FC-PC-N3-200-L1, 1m, RWD, China), a optic fiber commutator (R-FC-1x1, RWD, China), and a splitter branch patch cord (FCM-2xZF1.25(F), NA 0.37, Doric, Germany). The laser intensity during the behavioral experiments was 9-10 mW, and verified with a laser power meter (UT385, UNI-T, China) before the experiments. In trace fear conditioning (day 1), the rats received eight CS-US pairings. The CS-US pairing consisted of tones (2.5kHz, 20 s, 75 dB) and a footshock (0.7 mW, 1 s). CS and US are separated by 20 s trace interval. To investigate the role of CPn and COM neurons during the trace interval, the laser was illuminated during trace period in each trial. The intertrial interval of the fear conditioning was 180 ± 20 s. The next day, in the tone test (day 2), the rats received twenty presentations of tones (20 s, 2.5 kHz, 75 dB) in context B. During the tone test, to investigate the role of sustained increased activity in CPn and COM neurons during post-CS period, the laser was illuminated during a post-CS period (20 s) in each trial. The intertrial intervals of tone presentations were 90 ± 20 s.

For in vivo electrophysiology recordings after TPP or PBS infusion, TPP or PBS was infused 30 minutes before the conditioning (**Fig. 2a, 6a**) or before the tone test (**Fig. S3d-g**). In trace fear conditioning (day 1), three habituation tones were presented first to test whether PL neurons respond to the CS before CS and US are associated. These tone-only trials were followed by eight CS-US pairings. The tone test was the same as the trace fear conditioning with optogenetic silencing experiment described above.

#### Operant delayed match to position (DMTP)

To evaluate performance in a working memory task, we employed an operant delayed match-to-position (DMTP) task. The apparatus featured two visual stimuli (white squares) displayed above two corresponding nose-poke holes. Rewards were dispensed through a lick port located on the opposite side of the visual stimulus. The reward delivery was signaled by light illumination. We used cherry-flavored Kool-Aid (KHC, Chicago) at a concentration of 0.4 g/ml as reward. A session, each lasting one hour, progressed through six stages: habituation, pretraining, match, 1 s DMTP (training session), 4 s DMTP (challenging and test sessions). During the habituation stage, animals underwent reward magazine training. They received 2.5 μl of reward at 15 s intervals, with each delivery accompanied by reward light illumination. The light extinguished when the animal licked the reward port. The following day, pretraining began with the presentation of a single visual stimulus. Animals received a 2.5 μl reward upon poking the nose-poke hole beneath the stimulus. This phase, lasting one hour, incorporated a 15 s inter-trial interval (ITI). Daily training continued until animals consistently achieved at least 30 trials per hour. To introduce the match-to-position sequence, we initiated the match phase. This stage began with a single visual stimulus (sample) presentation, followed by a 2.5 μl reward. Subsequently, the visual stimulus reappeared at the same location (choice), and a larger reward (6 μl) was provided. Incorrect responses, where the animal poked the opposite nose-poke, resulted in house light illumination for 5 s, followed by a 20 s ITI. Training continued until animals consistently performed 30 trials per hour with a 70% success rate. The 1s DMTP phase was the same as training session, built upon the match phase with two key modifications. First, we introduced a 1 s delay between reward light illumination and reward availability during the sample reward (2.5 μl) delivery. Second, we presented two visual stimuli at the choice stage instead of one. As before, animals were trained to achieve 30 trials per hour with a 70% success rate. In the 4 s DMTP phase, we extended the delay to 4 s. Animals were considered ready for experimental testing when they reached 30 trials per hour with a 70% success rate at this delay interval. In the last phase, test session, animals performed 4 s DMTP for two days, then received PBS or TPP 30 minutes before 4 s DMTP the next day and continue to perform 4 s DMTP for two additional days.

### Unit recordings and data processing

One week after drive implantation, the rats were allowed to acclimate to the laboratory for 15 minutes before trace fear conditioning. The 32-channel RHD recording headstage (C3314, Intan, USA) was connected to the Omnetics connector of the shuttle drive. PFC activity was recorded using Open Ephys software (Open Ephys, Portugal) and an RHD interface board (RHD USB interface board, Intan, USA). The headstage and RHD interface board were connected with a thin serial peripheral interface (SPI) cable (C3213, Intan, USA). All in vivo recordings were performed at a sampling rate of 30 kHz. A bandpass filter from 500 to 5000 Hz was used to remove background noises. The recording session was 34 minutes during the fear conditioning and 42 minutes during the tone test. The behavior of rats was recorded by a CCD camera. The in vivo recording and tone presentations were synchronized using Bonsai software (Bonsai Foundation). After finishing the in vivo recording, a 30 μA current was applied to each stereotrode for 8 s to locate the position of the tip of stereotrodes bundle. Brains were extracted and post-fixed for 24 h in the 4% PFA. The coronal prefrontal cortex (300 μm thick) was prepared using a vibratome and imaged with a fluorescence microscope (IX53, OLYMPUS, Japan).

For spike sorting, we modified a Matlab software package published in github (https://gith ub.com/nghorbani/NeuralDataAnalysis). Before in vivo recording data were subject to spike sorting, we concatenated the in vivo recordings of each trial from -20 s to +60 s when the onset time of tone in each trial was zero. Spike waveforms were detected by amplitude thresholding. The standard deviation (σ) of noise was calculated as a median of absolute deviation (MAD) of filterd signal divided by 0.6754 ^28^, and the signal whose amplitude was larger than 5 σ was classified as a spike. The 12 and 30 data points before and after the peak of each spike were collected into a spike wavelet matrix. we refined the list of automatically detected spike waveforms by excluding spikelet such that the valley/peak amplitude ratio was greater than 0.8 or the peak amplitude was greater than 1000 μV. Next, the spikelet waveform matrix (time × spikes × channel) was suject to principal component analysis (PCA) to extract three principal features of spikelets per a channel. The extracted features were plotted on the coordinate of six principal components (three principal components × two channels), to which a Gaussian mixture model was fitted for clustering using split-and-merge expectation-maximization (SMEM) algorithm ^29^. For further manual sorting, we used MatClust software (Mattias Karlsson, 2022). Spikes whose width greater than 200 μs and firing rate less than 15 Hz were classified as putative pyramidal neurons, and used for further analysis (**Fig. S6b-e**).

To select cells displaying distinctly high activity during CS and post-CS period (20 - 40 s and 40 - 60 s in **Fig. 2c**, respectively), we calculated z scores for firing frequencies and interspike intervals (ISIs), denoted as zf and zisi, respectively, with reference to their baseline (pre-CS) values. For calculation of zisi, we adopted a test for comparing two Poisson means (Krishnamoorthy and Thomson 2004), because the distribution of inter-spike intervals (ISIs) of randomly firing neurons follows an exponential distribution. The zisi during CS and post-CS periods *vs*. baseline was calculated as follows:

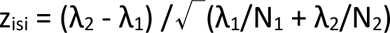

where λ1 is the mean of ISIs during baseline; λ2 is the mean of ISIs during CS or Post-CS; N is the sample size of each period. Cells satisfying one of two following criteria during CS or post-CS period were classified as CS and PA cells, respectively: (1) zf > 1.96 (p < 0.05); (2) zisi > 1.96 (p < 0.05) and zf > 1.28 (p < 0.1).

### Whole-cell patch recording

Brain slices were prepared from Sprague Dawley rats (21-28 days). Rats was sacrificed after being anesthetized with isoflurane, and the brain was immediately removed and chilled in ice-cold low-calcium artificial CSF (aCSF) containing the following (in mM): 110 choline chloride, 10 glucose, 25 NaHCO3, 2.5 KCl, 1.25 NaH2PO4, 2 Na-pyruvate, 3 ascorbate, 0.5 CaCl2 and 7 MgCl2 with pH 7.4 adjusted by saturating with carbogen (95% O2, 5% CO2). The coronal prefrontal cortex (300 μm thick) and visual cortex (300 μm thick) were prepared using a vibratome (VT1200S, Leica, Germany). Slices were recovered at 36°C for 30 minutes and after that maintained at room temperature with standard aCSF containing the following (in mM): 125 NaCl, 10 glucose, 25 NaHCO3, 2.5 KCl, 1.25 NaH2PO4, 2 Na-pyruvate, 3 ascorbate, 1.3 CaCl2 and 1 MgCl2 with pH 7.4 adjusted by saturating with carbogen. For recording, slices were transferred to a recording chamber and circulated with standard aCSF using a peristaltic pump (MINIPLUS3, Gilson, USA).

Whole-cell current clamp and voltage clamp recordings were performed at 32°C. These cells were visualized using an upright microscope mounted with differential interference contrast optics (BX61WI, Olympus, Japan). Electrophysiological recordings were made by the whole cell clamp technique with an EPC-10 amplifier (HEKA, Germany) or Integrated Patch Clamp amplifier (SUTTER instrument, USA) at a sampling rate of 10 kHz. Recording pipettes (3-5 MΩ) were filled with intracellular solution contained (in mM): 130 K-gluconate, 8 KCl, 20 HEPES, 4 MgATP, 0.3 NaGTP, 1 MgCl2 and 0.1 EGTA with the pH adjusted to 7.3 with KOH. EPSCs were recorded from the PL in a whole cell configuration at a holding potential of -65 mV. Series resistance (Rs) was regularly checked by applying a short hyperpolarization (20 ms, 2 mV). The data in which the Rs exceeded 20 MΩ or changed (>20%) during the experiment were discarded. For electrical stimulation, we filled a glass pipettes (2-3 MΩ) with aCSF and delivered monophasic stimulus pulses (2-15 mV, 100 μs/pulse) using a stimulus isolator (A360, WPI, USA). For L2/3 stimulation, the stimulation pipette was placed in L2/3 within 300 μm from the somata of a postsynaptic L5 cell.

For PTP induction, stimulation was adjusted to evoke baseline current amplitudes of 100 pA. For baseline EPSC recording, EPSCs were evoked by paired pulses (Inter-spike Intervals, 50 ms) every 5 s 20 times. PTP was induced by stimulation mimicking the firing patterns of an exemplar CS responsive L2/3 pyramidal neuron during CS period, as derived from in vivo electrophysiology recording (**Fig. 5m**). After the patterned stimulation, paired pulse stimulations were resumed for 50 times. Series resistance compensation, bridge balance, or liquid junction potential correction was not applied. The mNCX blocker, tetraphenylphosphonium chloride (TPP), was purchased from Sigma (218790, Sigma, USA).

### In vivo Ca^2+^ imaging and data processing

Three weeks after virus injection, rats were subject to trace fear conditioning. The next day, the rats were allowed to acclimate to the experimental room for about 15 minutes, and then the Miniscope (V4; Open Ephys, Portugal) was docked to the baseplate to record calcium signal. Calcium signals were recorded using open-source Miniscope DAQ software (Aharoni Lab, UCLA) and Miniscope. The Miniscope and DAQ were connected with a flexible coax cable, and the calcium signal was transmitted to the data acquisition system (DAQ) through the cable and saved on a personal computer. The frame rate, LED power, gain, and focus were also controlled using the Miniscope DAQ software. The calcium activity was recorded at 25 frames per second at a resolution of 600 **×** 600 pixels (LED intensity 2-15; gain 2.5-3.5). The behavior of rats was recorded by a CCD camera. The calcium imaging and tone presentations were synchronized by Bonsai software (Bonsai Foundation). To reduce a decrease in fluorescence intensity due to photobleaching, calcium signal was not recorded in the entire behavioral experiment, but was recorded separately for each trial. Calcium signal recording was started 30 s before the onset of the cue and was recorded for 110 s. To confirm the expression site of GCaMP6f and the location of GRIN lens implantation, brains were extracted and post-fixed for 24 h in the 4% PFA after finishing experiments. The coronal sections of prefrontal cortex (300 μm thick) were prepared using a vibratome and imaged with a fluorescence microscope.

The calcium imaging videos were down-sampled from 600×600 pixels to 300×300 pixels using ImageJ software (NIH, USA). The down-sampled calcium imaging videos over five tone trials were concatenated using ImageJ software and saved as one TIF file. The processed calcium imaging data were motion-corrected using NoRMCorre^30^ implemented in MATLAB. After down-sampling and motion correction, we processed the image data using a constrained non-negative matrix factorization algorithm optimized for microendoscopic data (CNMF-E)^16^. The data was filtered with a gaussian kernel with a width of 4 pixels (gSig), and neurons were constrained to a diameter of 15 pixels (gSiz). A ring model (bg_model) was used for the background with a radius of 18 (ring_radius). Neurons with spatial overlap greater than 0.2 (merge_thr) and centroid distance less than 10 pixels (dmin) were merged. Only ROIs with a minimum peak-to-noise ratio of 15 (min_pnr) and minimum spike size of 5 (smin) were extracted. The foopsi deconvolution method by CNMF-E was employed in all datasets. Not all ROIs classified by CNMF-E could be considered as neurons and some might be dendrites or background fluctuations. Therefore, all neurons were visually inspected, and those that displayed non-neuronal morphology were removed. Further data analysis was performed using the inferred spiking activity (neuron.S). After converting a non-zero value to 1 in the inferred spiking activity value (binarized spike activity), first five trials of tone test were averaged to reduce trial-to-trial variability of neuron activity pattern.

### Statistics

Data were analyzed with Igor Pro 7 (Wasvemetrics, USA), SPSS (IBM, USA), Prism (GraphPad, USA), MATLAB (Mathworks, USA), R (https://www.R-project.org/) and RStudio (http://www.rstudio.com). The sample size per group and statistical tests are mentioned in the respective figure legends. For freezing behavior, we used repeated measures two-way ANOVA followed by Tukey post hoc analysis. Post hoc comparisons with Bonferroni correction were conducted to investigate differences between groups at each time point. The difference in mean z-score between PBS and TPP injected groups during CS1, CS2 and Trace (or post-CS) periods was tested with a linear mixed effects (LME) model. For the LME model, time and drug treatment were set as fixed effects, and individual animals and neurons as random effects to account for repeated measures and clustering by animals^31^. The “nlme” package of R language was used for fitting an LME model^32^, and the “emmeans” package was used for pairwise comparisons^33^. The Wilcoxon matched-pairs signed rank test was used for two paired groups. A chi-square test was used to compare the proportions of neuronal activity patterns. The level of statistical significances was indicated as follows: n.s., not significant (p > 0.05), *P < 0.05, **p <0.01, ***p<0.001, ****p<0.0001. All data in the figures are presented as the mean ± standard error of mean (SEM).

