## supplementary figures for "Prelimbic corticopontine neurons gate extinction learning"

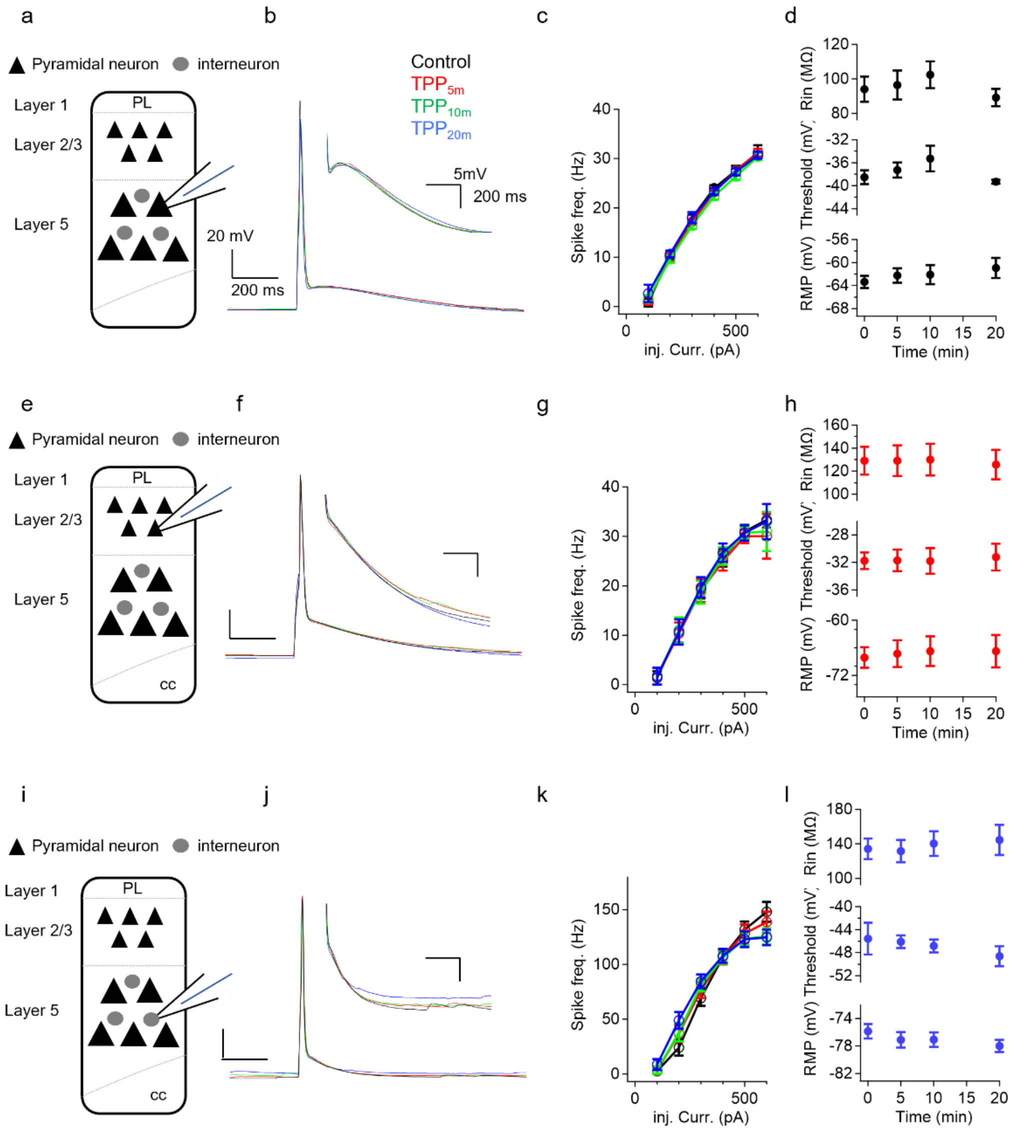


**Fig. S1. TPP does not affect intrinsic electrical properties of PL neurons. a-d,** TPP effects on L5 pyramidal neurons (PNs) in the PL area. After depolarization (ADP, *b*), firing frequency-current injection relationship (F-I curve, *c*), Input resistance ($R_{in}$, *d*), Resting membrane potential (RMP, *d*) and action potential threshold (AP threshold, *d*) were monitored in L5 PNs over time after TPP application. **a,** Scheme of experiments show the location of patch clamp pipette. **b,** Representative traces of APs and ADPs (*inset*) over time after TPP application (control = black, red = 5 min, green = 10 min blue = 20 min after TPP application). **c,** Averaged F-I curves of L5 PNs (n = 5). Same color codes as in *b*. **d,** $R_{in}$, RMP and AP threshold were not affected by TPP application (n = 7 for all). **e-h,** ADP, F-I curve, $R_{in}$, RMP and AP threshold in L2/3 PNs over time after TPP application. Control = black, red = 5 min, green = 10 min, blue = 20 min after TPP. Neither of ADP (*f*), F-I relationship (*g*), $R_{in}$ (*h*, upper), RMP (*h*, middle) and AP threshold (*h*, lower) was affected by TPP application (n = 6 for all). **i-l,** ADP, F-I curve, $R_{in}$, RMP and AP threshold in L5 INs over time after TPP application. Same color codes as in *b*. (n = 13 for all). All data are mean ± S.E.M.

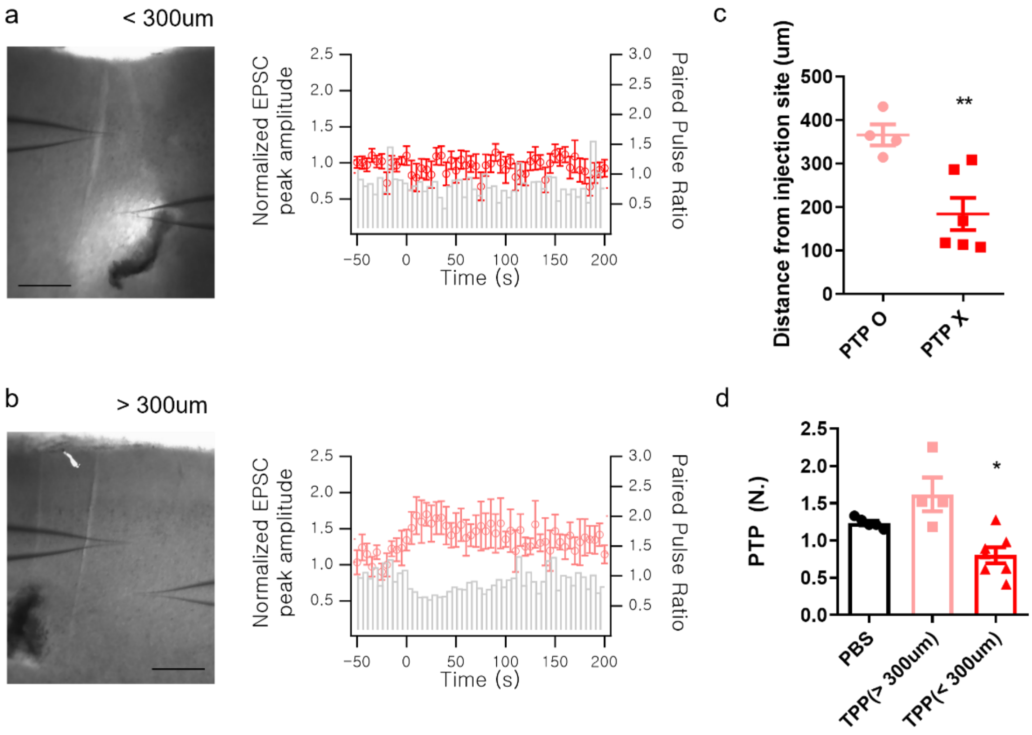


**Fig. S2. PTP was not induced within the area 300 μm from the injection site, but was induced outside the area. a,** PTP induction at L2/3-CPn synapses within the area 300 μm from the TPP injection site. *Left*, Representative image of PTP induction experiment within the area 300 μm from the TPP injection site showing the location of patch clamp pipette, monopolar stimulation glass electrode and PBS injection site. Scale bar, 200 μm. *Right*, Time courses of baseline-normalized EPSC amplitudes before and after tetanic simulation (TS, 25 hz for 5 s) applied at t = 0. PTP was not induced within the area 300 μm from the injection site (n = 7). **b,** PTP induction at L2/3-CPn synapses outside the 300 μm area. *Left*, Representative image of PTP induction experiment at outside the 300 μm area. Scale bar, 200 μm. *Right*, Time courses of baseline-normalized EPSC amplitudes before and after TS applied at t = 0. PTP was induced outside the area (n = 4). **c,** Summary scattered plot of distances from injection site *vs*. occurrence of PTP (P = 0.0095 for PTP o vs PTP x). **d,** Summary bar graphs for normalized PTP. PTP was not induced within the area < 300 μm from the TPP injection site, while PTP was induced outside the area. For PTP within (n=7) vs. outside (n=4) the 300 μm area, P = 0.0121. Mean ± S.E.M., *P < 0.05, **P < 0.01, Mann-Whitney test.

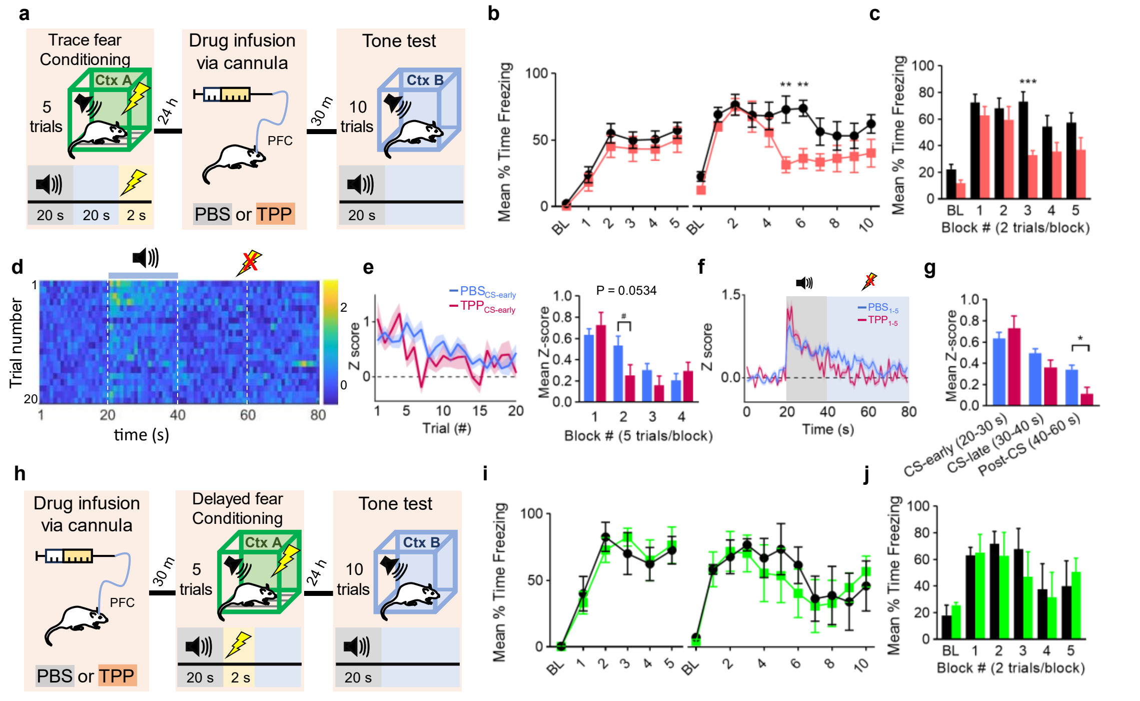


**Fig. S3. a-c, Injection of TPP into the PL before the tone test on D2 accelerated trace fear extinction similar to TPP injection before trace fear conditioning (TFC). a,** Schematic of experiment procedure. PBS or TPP was infused into the PL 30 minutes before tone test (day 2). **b,** Time courses of freezing behavior during trace fear conditioning (*left*, day 1) and the tone test (*right*, day 2) in PBS and TPP injected animals (PBS, n=9; TPP, n=7 rats). RM two-way ANOVA (PBS vs. TPP): for conditioning (day 1), $F_{1,14}$ = 0.794, P = 0.388; for tone test (day 2), $F_{1,14}$ = 11.140, P = 0.005. **c,** Mean freezing ratios of PBS and TPP-injected animals as a function of two trial blocks during tone test. RM two-way ANOVA (PBS *vs*. TPP): $F_{1,14}$ = 12.622, P = 0.003. **d-g, TPP injection into the PL before the tone test resulted in accelerated trace fear extinction, similar to the TPP injection before conditioning. d,** Time course of mean CS cell activities for 20 trials of the tone test. For each trial of the tone test, time series of 1 s binned z-scores were averaged over CS cells (D2; TPP, n = 21/77 neurons from 5 rats). **e,** *Left,* Trial-dependent changes of CS cell activity during first 10 s (CS-early) and late 10 s (CS-late) of tone presentation in PBS and TPP groups. *Right,* Mean z-scores of CS cells during CS-early period (20-30 s) across five trial blocks (5 trials/block). LME models (TPP *vs*. PBS): $F_{1,12}$ = 0.480, P = 0.0534, for trial block 2. **f,** For TPP group, z-scores of first five trials shown in *d* were averaged and displayed as a function of time. **g,** Mean z-scores of CS cells during three epochs (CS-early, CS-late and post-CS period) of first five trials of the tone test in PBS and TPP animals. LME models of 1 s binned CS cell activities (PBS vs TPP): for post-CS period (40-60 s), $F_{1,12}$ = 8.467, P = 0.0131. The PBS data in *e-g* were reproduced from Fig. 2 for comparison. **h-j, TPP injection to the PL did not affect delayed fear conditioning (DFC) on fear extinction. h,**

Schematic of DFC experimental procedure. PBS or TPP was infused into the PL 30 min before DFC on day 1. **i,** Time courses of freezing ratio of PBS (n=4) and TPP (n=4) animals during delay fear conditioning (*left*, day 1) and tone test (*right*, day 2). RM two-way ANOVA (PBS *vs*. TPP): for DFC, $F_{1,6}$ = 1.162, P = 0.981; for tone test, $F_{1,14}$ = 444.290, P = 0.694. **j,** Mean freezing ratio of PBS and TPP groups during tone test, presented as means of two trial blocks. RM two-way ANOVA (PBS *vs*. TPP): $F_{1,6}$ = 70.232, P = 0.874. Post hoc comparison was done using Bonferroni correction to determine the statistical difference between groups at each time point. All data are mean ± S.E.M., Shaded areas represent S.E.M., *P < 0.05, **P <0.01.

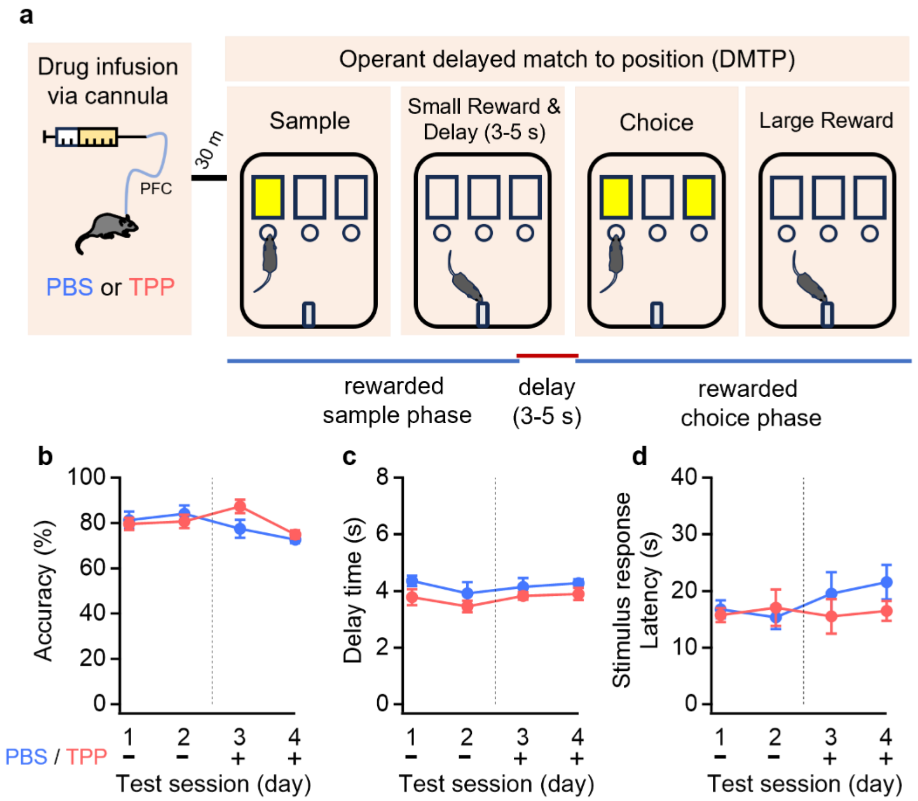


**Fig. S4. TPP injection into the PL did not affect performance on the working memory task. a** Schematic of operant delayed-match-to-position (DMP) task. Animals were subject to a test session of the DMP task 30 min after drug injection into PL. Each trial consists of a rewarded sample phase and a rewarded choice phase, separated by a delay of 1 s (training session) or 3-5 s (challenging session). If the performance accuracy reached 70% for 30 trials per an hour in the challenging session, the test session, consisted of 30 trials, started on the following day. At 30 min prior to the test session, PBS or TPP was infused into the PL. Performance in the DMP task did not differ between the two groups **b,** Performance accuracy before and after PL injection of PBS (n = 5; blue) or TPP (n = 6; red circles) during test sessions over four days. PBS (blue circles) or TPP (red circles) was injected on the 3rd day (A vertical dashed line). **c,** Delay time between sample and choice phase of PBS and TPP injected groups. **d,** stimulus response latency of PBS and TPP injected groups. All data are mean ± S.E.M.

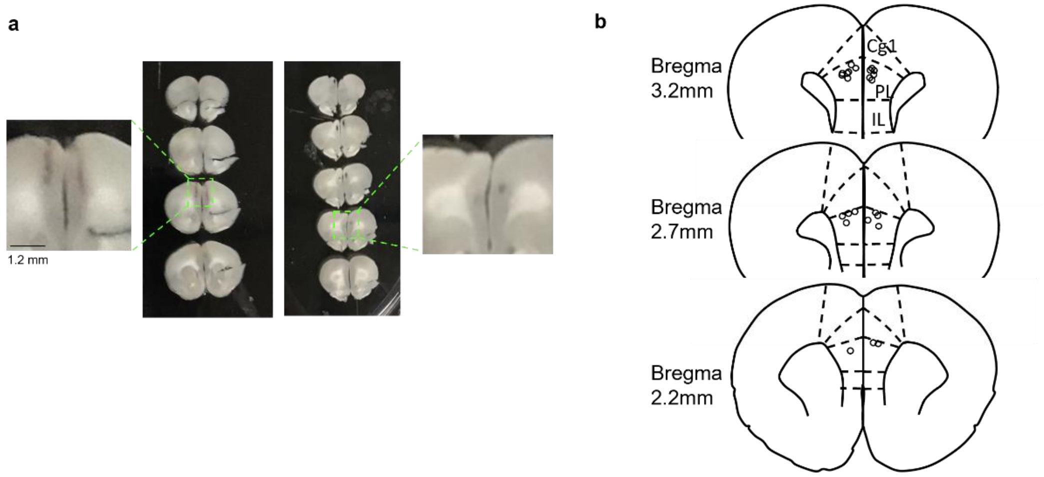


**Fig. S5. Bilateral infusion cannula implantation into the PFC PL area. a,** Representative images of coronal slices of the brain implanted with bilateral cannula. **b,** locations of bilateral infusion cannula of all rats used in the present study.

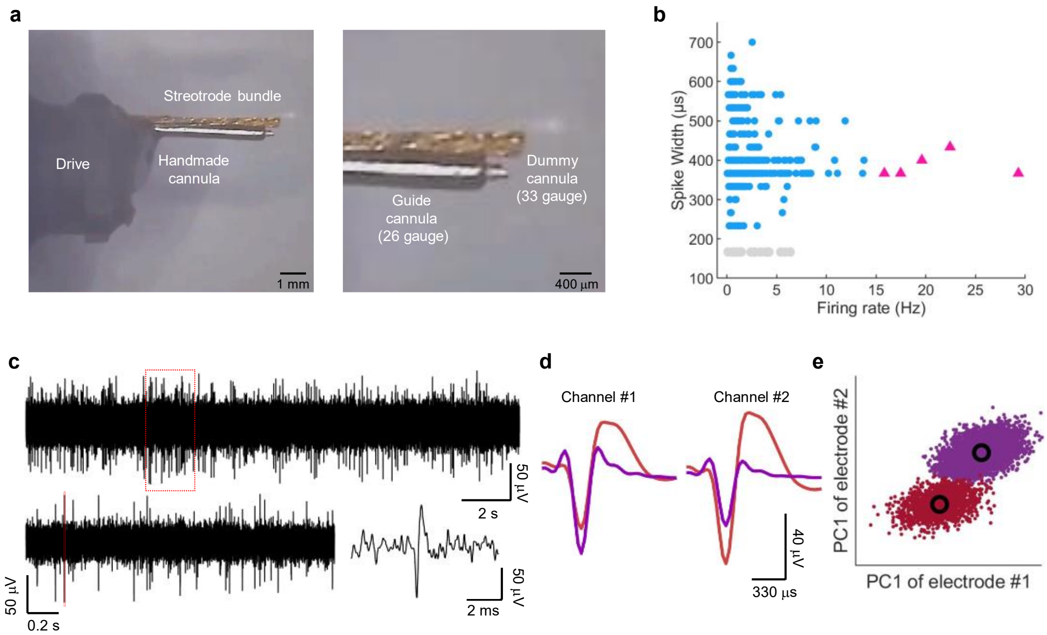


**Fig. S6. In vivo electrophysiological recording. a,** Modified shuttle drive (Voigts et al. ,2019). *Left,* Representative image showing a stereotrode bundle and handmade cannula in close proximity. Scale bar, 1 mm. *Right,* Magnified image showing a guide cannula (26 gauge), into which a dummy cannula (33 gauge, same size as infusion cannula) is inserted. The dummy and infusion cannula were set slightly shorter than the stereotrode bundle to ensure an optimal drug delivery at the recording site. Scale bar, 400 μm. **b,** Scatter plot of sorted spikes on the plane of spike width vs. firing rate for all recoding (n = 381). Sorted spikes were classified into putative pyramidal neurons (blue dots) and interneurons (red triangle and gray dots) based on their waveform durations and firing rates. **c,** Representative traces recorded *in vivo* using a stereotrode at different time scales. The electrophysiological signal was recorded at 30 KHz and band-pass filtered from 0.5 to 5 kHz. The boxed region of upper trace was expanded on time and shown in lower trace. **d-e,** Representative unit spike waveforms (*d*, called spike wavelet, 42 data points/wavelet) recorded during TFC. Scale bar, 20 μV, 330 μs. Such Spike wavelets were collected into a matrix of three dimensions (spike x time x channel). The wavelet matrix was then subject to PCA, yielding three principal components at each of two electrodes of a stereotrode. The two spike wavelets shown in *d* made two separate clusters on the plane of two first principal components from two channels (*e*). Spikes were sorted by fitting a Gaussian mixture model to the extracted features on the space of six principal components, and clustering based on split and merge expectation maximization algorithm (Ueda et al., 2000).
